# Holistic Characterization of Tumor Monocyte-to-Macrophage Differentiation Integrates Distinct Immune Phenotypes in Kidney Cancer

**DOI:** 10.1101/2021.07.07.451502

**Authors:** Adriana M. Mujal, Alexis J. Combes, Arjun R. Rao, Mikhail Binnewies, Bushra Samad, Jessica Tsui, Alexandre Boissonnas, Joshua L. Pollack, Rafael J. Argüello, Megan K. Ruhland, Kevin C. Barry, Vincent Chan, Matthew F. Krummel

## Abstract

The tumor immune microenvironment (TIME) is commonly infiltrated by diverse collections of myeloid cells. Yet, the complexity of myeloid cell identity and plasticity has challenged efforts to define *bona fide* populations and determine their connections to T cell function and their relation to patient outcome. Here we leverage single-cell RNA-sequencing (scRNA-seq) analysis of several mouse and human tumors and find that monocyte-macrophage diversity is characterized by a combination of conserved lineage states as well as transcriptional programs accessed along the differentiation trajectory. Using mouse models, we also find that tumor monocyte-to-macrophage progression is profoundly tied to regulatory T cell (Treg) abundance. Importantly, in human kidney cancer, heterogeneity in macrophage accumulation and myeloid composition corresponded to variance in, not only Treg density, but also the quality of infiltrating CD8^+^ T cells. In this way, holistic analysis of monocyte-to-macrophage differentiation creates a framework for critically different immune states in kidney tumors.

## Introduction

A key component of most immune responses, including those to cancers, are mononuclear phagocyte cell populations, which share common features of phagocytosis, tissue repair, and immunoregulation but diverge in functional specialization. Conventional dendritic cells (cDCs) are positioned in tissues to initiate and sustain adaptive T cell responses^1, 2^ while macrophages (MFs) engage in high rates of phagocytosis and tissue remodeling^3–5^. Self-renewing tissue-resident macrophages are seeded during embryonic development^6–10^, while inflammatory stimuli prompt infiltration of adult hematopoietic stem cell-derived monocytes that give rise to tumor macrophages^11–14^. These monocyte-derived macrophages preferentially accumulate as tumors progress^15^ and may predominate in regulating the ongoing antitumor T cell response^16^.

Macrophages consist of numerous subset populations that have been identified across tissues^17–20^. Therapeutic blockade of key epigenetic and signaling pathways has demonstrated their amenability to transcriptional reprogramming^21–27^, but how phenotypic diversity arises still remains poorly understood. Recruited bloodborne monocytes exhibit plasticity in differentiation potential and can acquire features of macrophages and/or DCs depending on the inflammatory setting^12, 13, 17, 28–32^. In addition, early studies demonstrated that macrophage exposure to type 1- or type 2-associated cytokines induces “M1” or “M2” cellular programs, respectively, and a model was put forth in which myeloid cells are polarized to be pro- (“M1”) or anti- (“M2”) inflammatory^33–35^. Although this nomenclature was thereafter understood to require nuance to account for additional plasticity^36^, it remains undetermined if these binary programs are applicable to describe tumor macrophage differentiation *in vivo*.

Myeloid phenotypic diversity has also challenged efforts to utilize myeloid populations as biomarkers for patient treatment options and outcome. cDCs are critical for coordinating antitumor T cell immunity^37–42^ and higher cDC abundance is broadly associated with improved cancer patient survival, although additional TIME features may inform functionality^38, 39, 43^. In contrast, macrophages have largely been considered to be pro-tumoral^5, 44^ and monocytes have often been described as myeloid-derived suppressor cells (MDSCs)^45^. Yet, several studies have exhibited variability in the use of macrophages as a negative predictor of patient prognosis^46–49^, and increased levels of circulating monocytes were unexpectedly linked to patient responsiveness to immune checkpoint blockade (ICB)^50^. These contrary findings speak to the need for improved resolution of myeloid cell categorization and phenotype in order to dissect heterogenous responses amongst cancer patients.

We used scRNA-seq to uncover transcriptional heterogeneity amongst tumor-infiltrating myeloid cells and distinguished monocyte and macrophage lineage- and activation-induced programs shared between multiple mouse tumor models and human kidney cancer samples.

Monocyte differentiation is dynamically regulated, and we found that Treg density is one example of an immunoregulatory axis that can modulate macrophage density. Further comprehensive analysis of key myeloid populations revealed distinct network connections between different myeloid cell types and T cell subsets, including Tregs and effector T cells. This is consistent with an archetypal organization of immune systems in tumors — collections of cell types that move together as modules^51^ — and improved classification of patients such that we could identify those with effective antitumor T cell responses.

## Results

### Establishing Diversity of Differentiation and Environment-responsive Myeloid States in Mouse B16 Tumors

Subcutaneous implantation of B16 melanoma cells is a well-established mouse tumor model with abundant infiltration of monocytes, macrophages, and cDCs^38^. To study these cells along their differentiation trajectories, we used conventional markers to sort bulk myeloid populations, along with reference sorted populations of Ly6C^+^ monocytes and two tumor-associated (TAM) populations, distinguished based on expression level of CD11c and MHC-II^38^ (**Fig 1A, S1A**), and then subjected all of these to scRNA-seq analysis.

**Figure 1.**
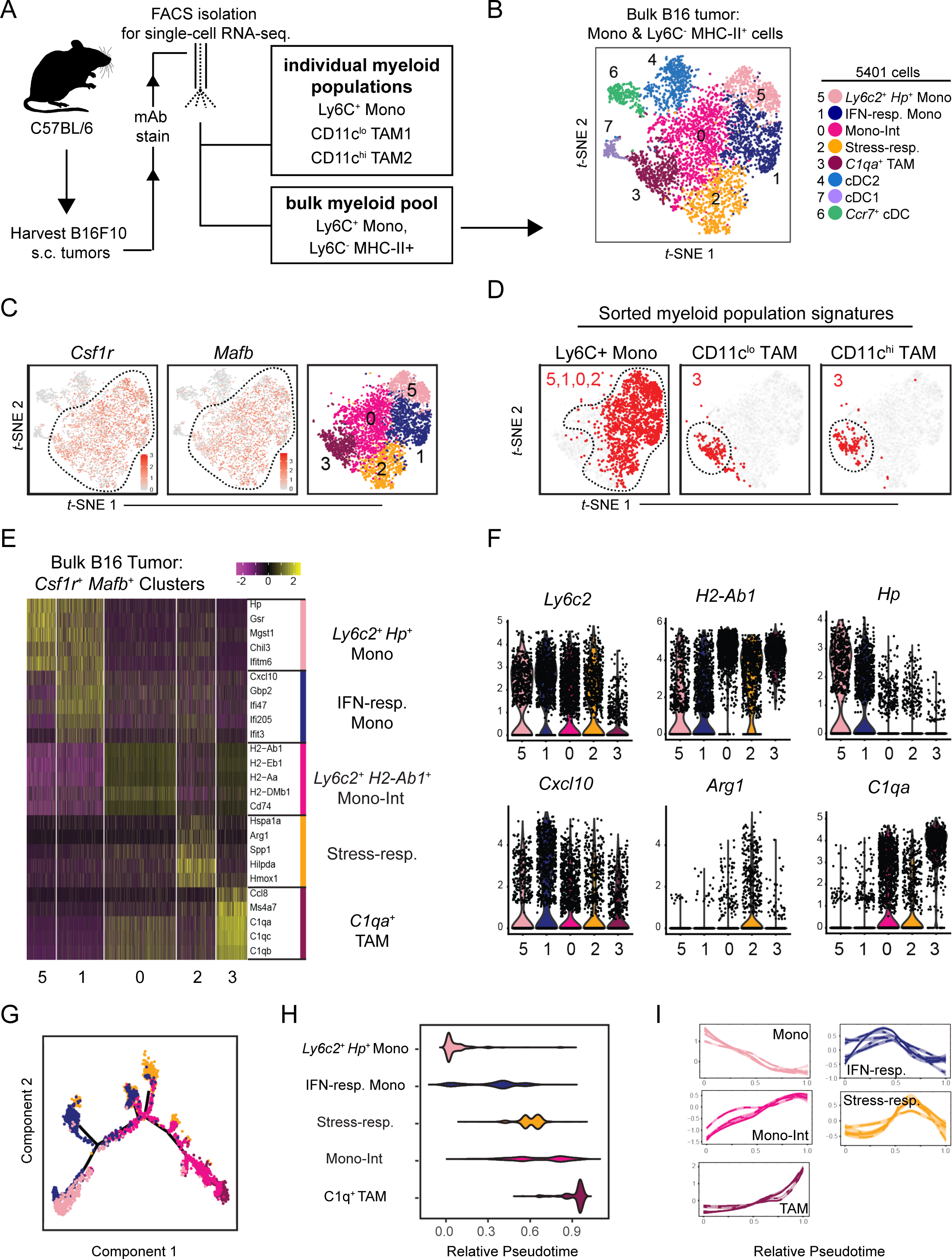
ScRNA-seq analysis of mouse B16 tumor myeloid cells maps transcriptional heterogeneity amongst monocytes and TAMs. **(A)** Schematic illustration of workflow for isolation of specified myeloid populations from B16 tumors subcutaneously implanted in wild-type C57Bl/6 mice. **(B)** *t*-SNE plot of graph-based clustering of Ly6C^+^ CD11b^+^ monocytes and Ly6C^-^ MHCII^+^ myeloid cells that were sorted and pooled from B16 tumors, and underwent scRNA-seq (**A**). **(C)** Expression of *Csf1r* (**left**) and *Mafb* (**middle**) on *t*-SNE plot of bulk myeloid cells (**B**), and display of selected *Csf1r*^+^ *Mafb*^+^ clusters (**right**). **(D)** Expression of gene signatures specific to Ly6C^+^ monocyte, CD11c^lo^ TAM1, or CD11c^hi^ TAM2 populations (**A, Fig. S1E**) displayed on *t*-SNE plot of *Csf1r*^+^ *Mafb*^+^ myeloid cells (**C**). Cells with top median of signature expression level labeled in red. **(E)** Heatmap displaying expression levels of top 5 differentially expressed (DE) genes between *Csf1r*^+^ *Mafb*^+^ cell clusters (**C**). Genes ranked by fold change. **(F)** Expression levels of selected genes amongst *Csf1r*^+^ *Mafb*^+^ cell clusters (**C**). **(G)** Differentiation trajectory model using Monocle analysis of cells from *Csf1r*^+^ *Mafb*^+^ clusters (**C**). Color coding corresponds to previous labels (**B**). **(H)** Graph of relative pseudotime values of *Csf1r*^+^ *Mafb*^+^ cluster cells (**C**) from Monocle analysis (**G**). **(I)** Expression levels of cluster-specific genes (**E**) over relative pseudotime (**H**). Each line corresponds to an individual gene.

Within the bulk myeloid population, *t-*SNE clustering yielded eight transcriptionally-distinct cell populations (**Fig. 1B, S1B-C**), including three *Flt3*^+^ *Kit*^+^ cDC populations (Clusters 4, 6, 7), which were marked by signatures specific to cDC1s, cDC2s, and conserved cDC activation programs (**Fig. S1D**, **Table S1**)^38, 39, 52^. The remaining myeloid cells (Clusters 0, 1, 2, 3, 5) broadly expressed *Csf1r* and *Mafb* (**Fig. 1C**), indicative of monocytes and macrophages. Having focused on the stimulatory capacity of cDC in previous work^38, 39, 53^, here we focused on the diversity of monocytes and macrophages as it related to the TIME.

To align transcriptional cell type categorization with flow cytometry analysis, we generated cell-type-specific gene signatures from the scRNA-seq analysis of the FACS-sorted monocytes and TAMs (**Fig. 1A**, **S1E**). When applied (**Fig. 1D**), these indicated that four *Csf1r*^+^ *Mafb*^+^ populations (Clusters 0, 1, 2, 5) expressed monocyte-specific genes, an unexpected heterogeneity. The four monocyte populations expressed *Ly6c2*, but varied in levels of other monocyte-associated genes (e.g., *Hp*, *Chil3*) and, as found in Cluster 0, also expressed TAM-associated genes (e.g. *H2-Ab1*, *C1qa*, *Ms4a7*) (**Fig. 1E-F, S1E**). Monocyte-like clusters were differentiated from one another by cellular activation programs. For example, Cluster 1 (“IFN-responsive”) was specifically enriched for interferon (IFN)-inducible genes such as *Cxcl10*, *Gbp2*, and IFIT-family members. Cluster 2 (“stress-responsive”) cells expressed *Arg1* and were enriched for cellular stress processes, including oxidative stress-responsive genes and heat-shock protein genes such as *Hmox1*, *Hspa1a*, *Hilpda*, *Bnip3*, *Ero1l*, and *Ndrg1* (**Fig. 1F**, **S1F**). In contrast to the heterogeneity observed amongst monocytes, signatures for both populations of TAMs localized within Cluster 3 (**Fig. 1D**).

We applied pseudotime analysis^54^ to generate a model of tumor monocyte-to-macrophage differentiation (**Fig. 1G-H, S1G**). This model placed Cluster 5 *Ly6c2*^+^ *Hp*^+^ monocytes and Cluster 3 *C1qa*^+^ TAMs at opposite ends of a linear trajectory consistent with expectations. Cluster 0 monocytes occupied the continuum between them and expressed a combination of both monocyte- and TAM-associated signatures such that we designated these cells “Intermediate monocytes” (“Mono-Int”). Kinetic analysis of cluster-enriched genes confirmed gradual downregulation of *Ly6c2*^+^ *Hp*^+^ monocyte-associated genes and up-regulation of “Mono-Int”- and TAM-associated genes along the pseudotime trajectory (**Fig. 1I**). This transcriptional model thus supported a framework of progressive monocyte-to-TAM differentiation, in which Ly6C down-regulation is paired with up-regulation of CD64, MHC-II, and F4/80^28^ (**Fig. S1H-I**). In contrast, IFN- and stress-responsive cells occupied distinct branches that diverged from the dominant differentiation trajectory at intermediate timepoints (**Fig. 1H-I, S1G**).

### ScRNA-seq Highlights Heterogeneous Acquisition of ‘Stress-’ and ‘Interferon-Responsive’ Cellular Programs during Tumor Monocyte-to-Macrophage Differentiation

To gain higher resolution on the differentiation trajectories within this lineage, we next performed cluster analysis on the sorted monocyte and TAM samples. Sorted monocytes expressed *Ly6c2* and contained clusters similar to those identified within the bulk myeloid cell sample (**Fig. 2A, S2A**). Cluster analysis of CD11c^lo^ and CD11c^hi^ TAMs, however, resolved diversity beyond the *C1qa*^+^ TAM signature (**Fig. 2B, S2B-C**) including clusters enriched for cell cycle-related genes, and an *Mgl2*^+^ TAM subset that expressed immune modulators such as *Ccl6*, *Il1b,* and *Retnla* as compared to the *C1qa*^+^ cluster which more highly expressed genes such as *Ms4a7*. Although these cells had not formed a distinct population in our original analysis of bulk myeloid cells (**Fig. 1**), we did retrospectively detect *Mgl2*^+^ cells in in that scRNA-seq data, as well as by flow cytometry (**Fig. S2D**). TAM-subset clusters were surprisingly also accompanied by an *Arg1*^+^ stress-responsive cluster akin to that found in the sorted monocytes (**Fig. 2B, S2B-C**). Indeed, re-clustering of the entire stress-responsive cluster from the bulk tumor myeloid sample revealed that this program was acquired by monocytes, “Mono-Int” and TAMs (**Fig. 2C, S2E**).

**Figure 2.**
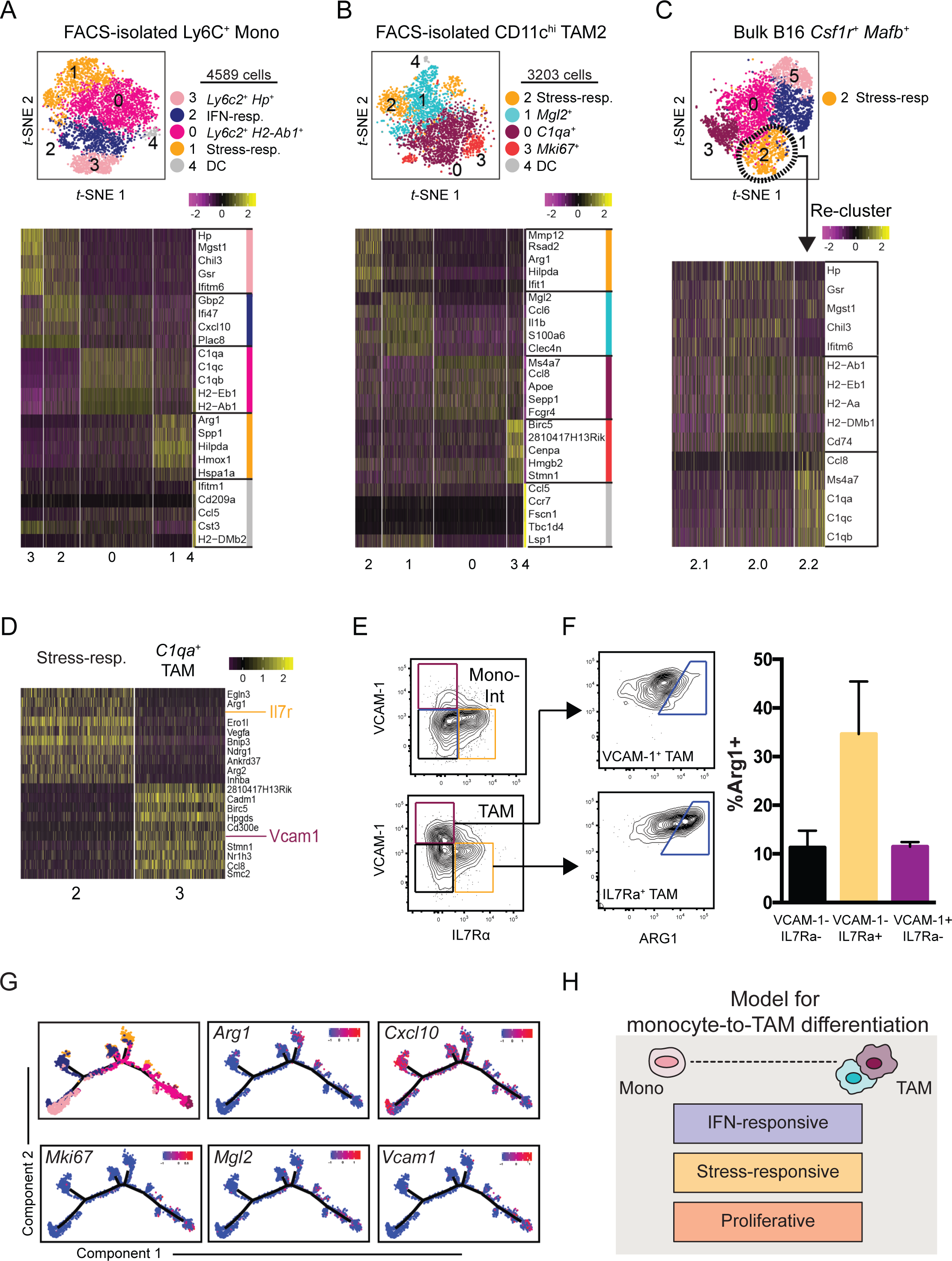
ScRNA-seq analysis highlights layering of microenvironment-induced programs during tumor monocyte-to-macrophage differentiation. **(A)** *t*-SNE plot of graph-based clustering (**top**) of Ly6C^+^ monocytes sorted from B16 tumors and processed for scRNA-seq (Fig. 1A), and heatmap displaying expression levels of top 5 DE genes between clusters **(bottom**) with genes ranked by fold change. **(B)** *t*-SNE plot and graph-based clustering (**top**) of CD11c^hi^ TAMs sorted from B16 tumors and processed for scRNA-seq (Fig 1A), and heatmap displaying expression levels of top 5 DE genes between clusters **(bottom**) with genes ranked by fold change. **(C)** Stress-responsive cells (Cluster 2) from bulk B16 myeloid cells (Fig. 1B) were selected for further clustering analysis (**top**). Heatmap of expression levels of monocyte- and macrophage-specific genes (Fig. 1E) by Cluster 2 sub-clusters (**bottom**). **(D)** Heatmap of DE gene expression levels between Cluster 2 and Cluster 3 of bulk tumor myeloid sample (Fig. 1B). Genes ranked by degree of exclusivity to a given cluster (min.pct1/min.pct2). **(E)** Expression levels of IL7R*α* and VCAM-1, as assessed by flow cytometry, of “Mono-Int” (Ly6C^+^ CD64^+^) (**top**) and TAMs (Ly6C^-^ F4/80^+^ CD64^+^) (**bottom**) from B16 tumors. **(F)** Example (**left**) and quantification (**right**) of intracellular ARG1 expression by VCAM-1^+^ (**top**) or IL7r*α*^+^ (**bottom**) TAMs from B16 tumors using flow cytometry. ARG1^+^ gating determined by isotype control. Data are representative of 2 independent experiments with 3-5 mice (mean ± SEM). **(G)** Expression levels of selected genes along differentiated trajectory generated by Monocle (Fig. 1G). **(H)** Schematic model of tumor monocyte-to-macrophage differentiation that integrates lineage-associated and microenvironmentally-induced transcriptional programs.

Segregated expression of stress-responsive genes and canonical TAM-associated genes suggested divergent transcriptional programs and we sought to determine if these populations could also be distinguished by flow cytometry. Differential gene expression analysis of the stress-responsive and *C1qa*^+^ TAM clusters from our bulk myeloid cell sample revealed cluster-specific expression of cell surface genes *Il7r* and *Vcam1*, respectively (**Fig. 2D**). Using the same gating as in **Fig. S1A**, we confirmed this split in both “Mono-Int” and TAMs (**Fig. 2E**) and we found enriched arginase 1 (ARG1) expression in both IL-7R*α*^+^ populations (**Fig. 2E-F, S2F**). As expected from the single-cell transcriptional analysis, VCAM1^+^ cells were more abundantly found within TAMs (**Fig. 2E-F, S2F**).

Together, this dissection of sorted cell populations lent support to a model in which monocytes and TAMs exist in a differentiation trajectory, along which cells can adopt specialized cellular programs (**Fig. 2G-H**). Some programs, such as those associated with *Mgl2*^+^ or *Vcam1*^+^ TAMs, selectively emerged later, in mature TAMs. Others, such as IFN-induced signaling or stress-responsiveness may be more universally accessible across differentiation stages. Interestingly, we detected populations of IFN-responsive monocytes in the peripheral blood of B16 tumor-bearing mice (**Fig. S2G-H**), perhaps suggesting that systemic IFN signaling, or other induction of this program, may define monocytes prior to tumor entry. In contrast, stress-responsive populations were not detected in the blood, suggesting that microenvironmental cues in the TIME likely induce this activation program locally. Further studies are warranted to explore if these programs directly influence monocyte differentiation processes or act as ‘layers’ that accessorize a canonical differentiation trajectory.

### Mouse Tumor Macrophage Subset Heterogeneity Does Not Reflect “M1/M2” Polarization

Macrophage exposure to type-1 or type-2 cytokines *in vitro* results in “M1” and “M2” transcriptional signatures that are often used to describe ‘pro-inflammatory,’ or ‘anti-inflammatory’ and wound healing processes, respectively^33–35^. To address whether “M1/M2” polarization was a useful construct to define tumor macrophage diversity *in vivo*, we tested how “M1” and “M2” gene signatures^55^ corresponded to tumor myeloid cell subsets profiled here.

Using correlation and clustering analyses (**Fig. 3A, S3A**), we found that, contrary to *in vitro* findings, tumor myeloid cells were marked by broad expression of both “M1”- and “M2”- associated genes, and we did not observe substantial correlation of gene expression within “M1” or “M2” gene groups across single cells. These data suggest that while tumor myeloid cells can express individual “M1” and “M2” genes, they rarely do so in any distinguishably consistent way during unperturbed tumor growth. Further, when a parallel sorting strategy was pursued to generate scRNA-seq analysis of tumor myeloid cells from the spontaneous mammary carcinoma MMTV-PyMT, we found that both share populations with the identical signatures defined in **Figure 1**, albeit in different proportions, and also show a lack of co-association between “M1” and “M2” signatures amongst the clusters (**Fig 3B-C, S3B-E**).

**Figure 3.**
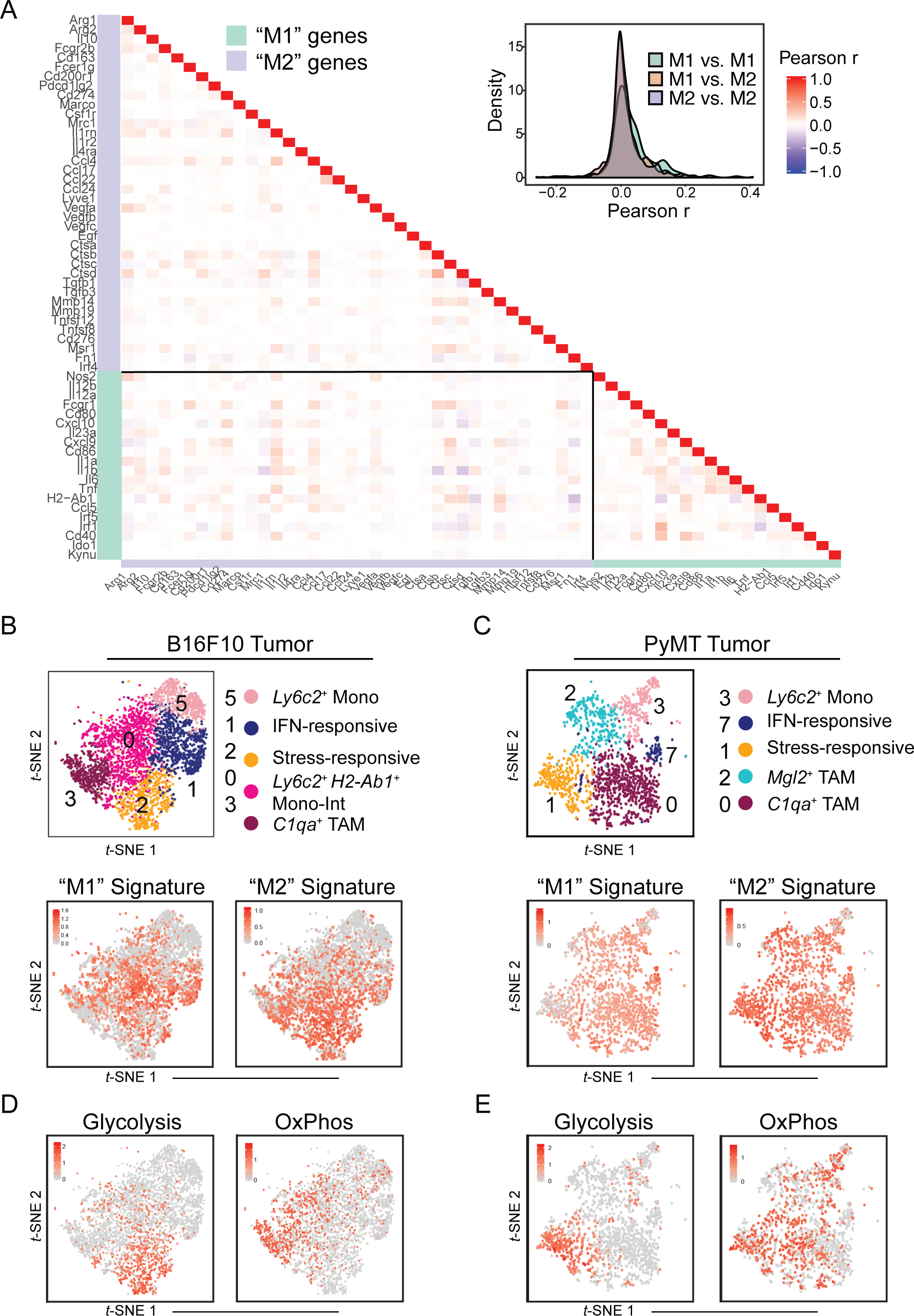
B16 and PyMT tumor monocyte/macrophage heterogeneity can be attributed to diversity in transcriptional and metabolic programs, but not “M1/M2” polarization. **(A)** Heatmap (**left**) and density plot (**right**) of Pearson r coefficient scores between “M1”- and “M2”-associated gene expression levels within *Csf1r*^+^ *Mafb*^+^ cells from B16 tumors (Fig. 1C). **(B)** *t*-SNE plot of *Csf1r*^+^ *Mafb*^+^ clusters from B16 tumors (**top;** Fig. 1C) with expression levels of “M1” **(bottom, left**) and “M2” (**bottom, right**) gene signatures (**A**) displayed. Cells with top median of signature expression level labeled in red. **(C)** *t*-SNE plot and graph-based clustering of *Csf1r*^+^ *Mafb*^+^ clusters from PyMT tumor myeloid cells that were sorted and processed for scRNA-seq (**top; Supplementary Fig. 3B**). Expression levels of “M1” **(bottom, left**) and “M2” (**bottom, right**) gene signatures (A) displayed. Cells with top 70 percentile of signature expression level labeled in red **(D)** Expression levels of glycolysis **(left**) and oxidative phosphorylation (“OxPhos”) (**right**) gene signatures (**Supplementary Fig. 3F**) displayed on *t*-SNE plot of *Csf1r*^+^ *Mafb*^+^ clusters from B16 tumors (Fig. 1C). Cells with top 70 percentile of signature expression level labeled in red **(E)** Expression levels of glycolysis **(left**) and oxidative phosphorylation (“OxPhos”) (**right**) gene signatures (**Supplementary Fig. 3F**) displayed on *t*-SNE plot of *Csf1r*^+^ *Mafb*^+^ clusters from PyMT tumors (**C**). Cells with top 70 percentile of signature expression level labeled in red

While myeloid cell populations appeared to be largely defined by differentiation stage and activation programs, we considered whether other core cellular features could help to further distinguish subsets across diverse microenvironments. It is now increasingly appreciated that metabolic reprogramming accompanies differentiation of immune cells, including macrophages^56^. Indeed, assessment of metabolism-related genes^57^ demonstrated that glycolysis-associated genes were specifically enriched in the stress-responsive cell cluster whereas genes pertaining to oxidative phosphorylation were specifically enriched in *C1qa*^+^ TAMs in two distinct mouse models (**Fig. 3D-E, S3F**). This suggests that these populations have additional important biological features in common—namely those coupled to distinct bioenergetic processes and demands.

Together, our data provides compelling evidence that “M1” and “M2” pathways have limited use in defining *in vivo* tumor myeloid cell differentiation and subset plasticity during normal tumor development. Rather, common microenvironmentally-induced programs and associated metabolic programs may yield greater insight in efforts to transcriptionally define and selectively target monocyte/TAM subsets.

### Human renal cell carcinoma-infiltrating monocytes and macrophages mirror murine populations

We then assessed how these mouse monocyte/macrophage transcriptional programs might compare to those from human kidney cancers, which are described to have substantial myeloid cell infiltration^58^. We performed scRNA-seq analysis on HLA-DR^dim/+^ Lin^-^ myeloid cells sorted from a renal cell carcinoma (RCC) sample (**Fig. 4A-B, S4A**). Signatures derived from previously described blood myeloid cell populations^59^ guided cluster identification and exclusion of cDC (**Fig. S4B**). Analysis of the *CSF1R*^+^ *MAFB*^+^ clusters revealed a heterogenous collection of monocytes and macrophages with varying levels of CD14 and CD16 (**Fig 4C-D**).

**Figure 4.**
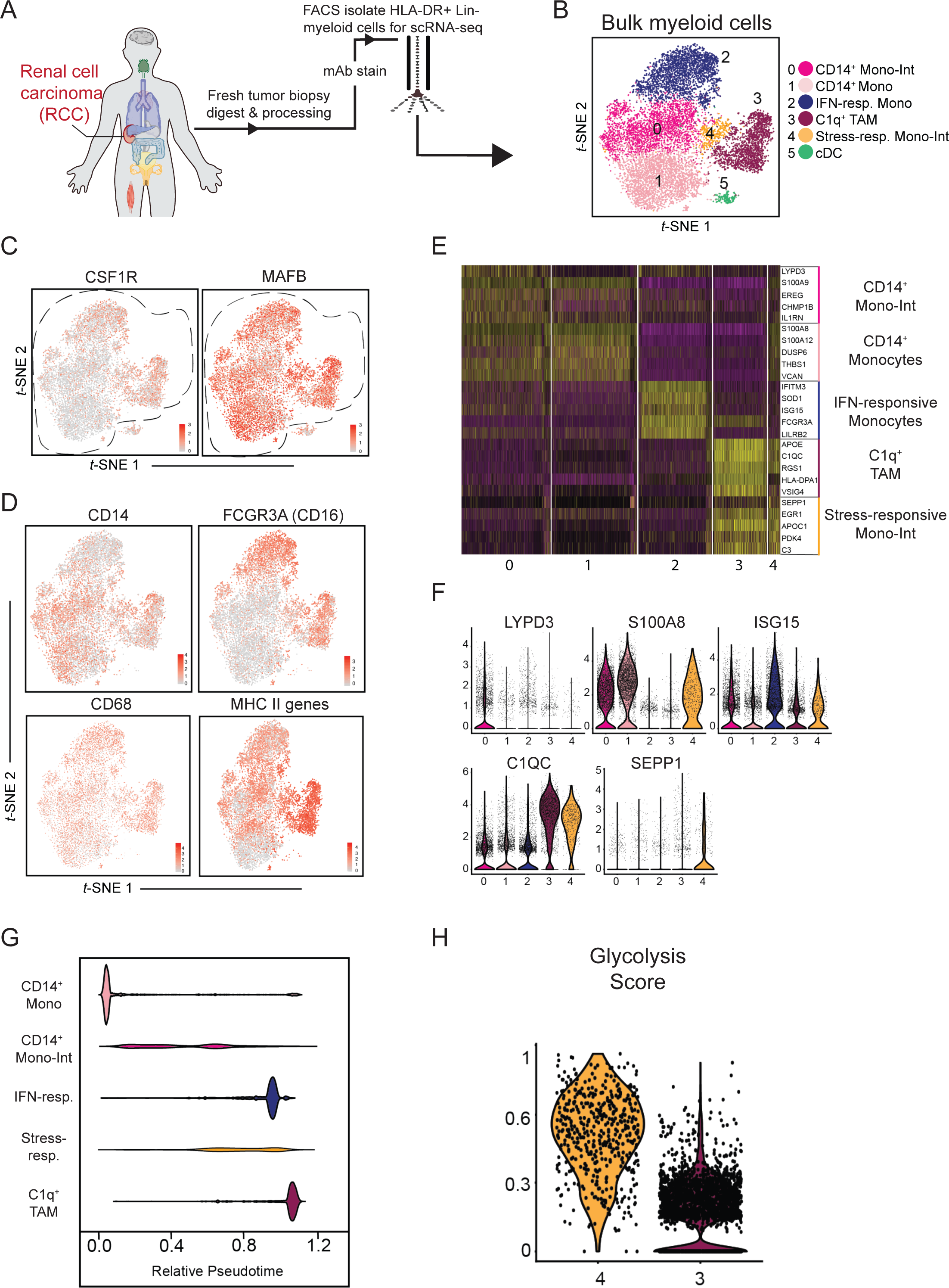
Human RCC and mouse tumor myeloid compartments exhibit shared transcriptional features. **(A)** Schematic of human RCC biopsy sample processed for scRNAseq analysis. **(B)** *t*-SNE plot of graph-based clustering of bulk myeloid (Lin^-^ HLA-DR^+^) cells sorted from human RCC biopsy sample (**A**). **(C)** Gene expression levels of *CSF1R* (**left**) and *MAFB* (**right**) displayed on *t*-SNE plot of human RCC-infiltrating myeloid cells (**B**). **(D)** Expression levels of selected genes (*CD14*, *FCGR3A*, *CD68*) or gene signature (MHC- II-associated genes) displayed on *t*-SNE plot of *CSF1R*^+^ *MAFB*^+^ clusters (**C**). **(E)** Heatmap of top 5 DE genes expressed by CSF1R^+^ MAFB^+^ clusters (**C**). Genes ranked by fold change. **(F)** Expression levels of selected genes by *CSF1R*^+^ *MAFB*^+^ cluster cells (**C**). **(G)** Differentiation trajectory model generated by Monocle analysis of *CSF1R*^+^ *MAFB*^+^ clusters (**C**). **(H)** Expression levels of glycolysis-associated gene signature by cells in stress-responsive (Cluster 4) and *C1Q*^+^ TAM (Cluster 3) cells (**B**).

As in mouse models, we detected early-stage *CD14*^+^ *S100A8*^+^ classical monocytes (Cluster 1) along with terminally-differentiated *C1QC*^+^ TAMs (Cluster 3) (**Fig. 4 E-F, Table S2**). Another population (Cluster 0) were *CD14*^+^ and differentially expressed *LYPD3* and MHC-II genes, consistent with intermediate differentiation of monocytes towards TAM (“Mono-Int”; **Fig. 4D-F**). A population of *CD16*^+^ non-classical monocytes (Cluster 2) also expressed IFN-stimulated genes and thus appear to functionally represent ‘IFN-responsive’ cells (**Fig. 4D-F, S4C**). Finally, we found that there were a mix of cells on the monocyte-macrophage trajectory that expressed the stress-responsive program identified in mice (**Fig. S4C**). Of these was a cluster transcriptionally similar to TAMs but marked by high expression of the antioxidant factor *SEPP1*^60–62^ (Cluster 4). When compared further to *C1Q*^+^ TAMs, this *SEPP1*^+^ cluster was less mature based on higher expression of monocytic markers (i.e. *S100A* genes) and lower expression of MHC-II-related genes (**Fig. 4E**, **S4D**). Pseudotime analysis of human myeloid cells recapitulated the alignment of stress- and IFN-responsive programs over the monocyte-to-macrophage trajectory (**Fig. 4G**), although in this RCC sample, IFN-responsive monocytes appeared more advanced in differentiation stage. As in the mouse samples, there was broad co-expression of “M1”- and “M2”-associated genes across the populations (**Fig. S4E**). Also, as in mice, there was a striking enrichment in a glycolytic signature^57^ within the stress-responsive (SEPP1^+^) cluster as compared to the C1q clusters, supporting that these cells were functional orthologs in the two species (**Fig 4H**). Altogether, these data confirm the limitation of “M1” and “M2” applicability in human tumors and illustrate the ability of other pathways to better define myeloid cell subsets *in vivo*.

### Treg depletion impairs monocyte-to-macrophage differentiation and elicits inflammatory monocyte programs

Myeloid cell density can vary across patients^58^, but how myeloid cell infiltration and differentiation is collectively regulated in human cancer is still not well understood. When we quantified myeloid cell populations in RCC patient biopsies using flow cytometry we found that the proportion of myeloid cells amongst live immune cells was increased in tumors of greater size and later stages (**Fig. S5A**). Closer examination revealed that the ratio of macrophages-to- monocytes was also specifically increased in more advanced tumors (**Fig. 5A, top**). This suggested that the balance between monocytes and macrophages is dynamically regulated and that tumor growth was tied to higher macrophage density.

**Figure 5.**
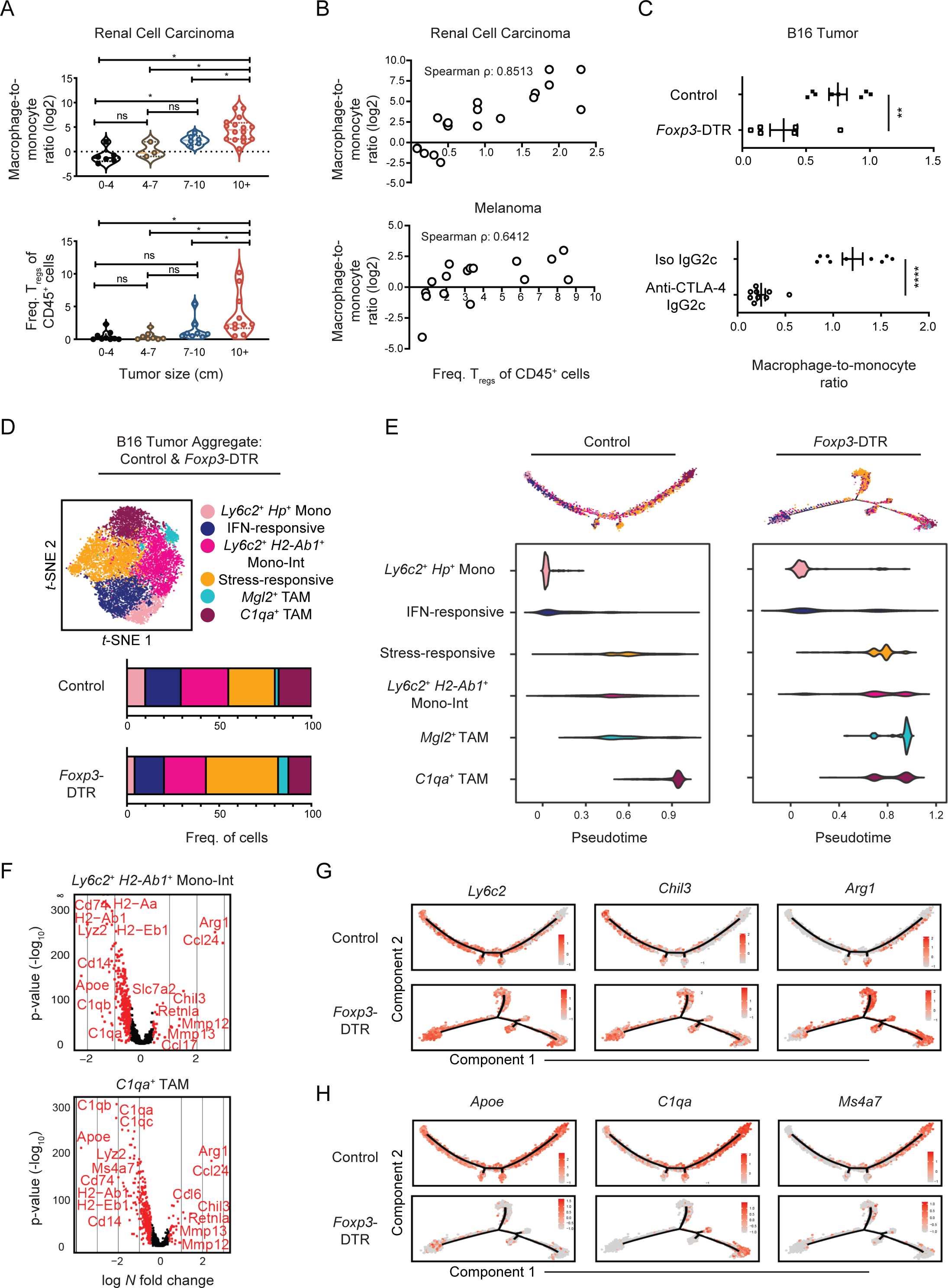
Immunosuppressive Treg cells promote tumor monocyte-to-macrophage differentiation. **(A)** Human RCC biopsies were measured and processed for flow cytometric analysis. The ratio of macrophage-to-monocyte (log2) cell numbers (**top**) and Treg frequency amongst CD45^+^ cells (**bottom**) were quantified. *p<0.05. **(B)** Dot plot and Spearman’s correlation coefficient of macrophage-to-monocyte cell number ratio (log2) and Treg frequency within CD45^+^ cells in human RCC (**top**) and melanoma (**bottom**) biopsies that were analyzed by flow cytometry. **(C)** Quantification of the ratio between macrophages (Ly6C^-^ F4/80^+^ CD64^+^) and monocytes (Ly6C^+^ CD11b^+^) cell number ratio in B16 tumors of DT-treated control and *Foxp3*-DTR mice (**top**), or of wild-type mice treated with depleting anti-CTLA-4 (IgG2c clone) or isotype antibody (**bottom**). Data is representative of 2 independent experiments. **p <0.01, ****p<0.0001. **(D)** *t*-SNE plot of graph-based clustering (**top**) of aggregated B16-infiltrating *Csf1r*^+^ *Mafb*^+^ cells from wildtype mice (Fig. 1), as well as DT-treated control and *Foxp3*-DTR mice (**Supplementary Fig. 5D**). Cell numbers in specified clusters were quantified (**bottom**). **(E)** Differentiation trajectory model generated from Monocle analysis (**top**) and relative pseudotime values (**bottom**) of *Csf1r*^+^ *Mafb*^+^ cluster cells from B16 tumors from DT- treated control (**left**) and *FoxP3*-DTR mice (**right**). **(F)** Volcano plot displaying DE genes between B16 tumor “Mono-Int” (**top**) and *C1qa*^+^ TAM (**bottom**) cluster cells from DT-treated control and *FoxP3*-DTR mice (**D**). Genes with > 0.4 log-fold changes and an adjusted p value of 0.05 (based on Bonferroni correction) are highlighted in red. Genes of interest labeled. **(G)** Expression of selected monocyte-associated genes displayed on the differentiation trajectory (**E**) of control (**top**) or *Foxp3*-DTR (**bottom**) B16 tumor-infiltrating *Csf1r*^+^ *Mafb*^+^ cells. **(H)** Expression of selected macrophage-associated genes displayed on the differentiation trajectory (**E**) of control (**top**) or *Foxp3*-DTR (**bottom**) B16 tumor-infiltrating *Csf1r*^+^ *Mafb*^+^ cells.

We thus sought other immunosuppressive parameters that might work in concert with increased macrophage accumulation. Tregs are a potent immunosuppressive force in the TIME and ablation can result in tumor clearance^39, 63^. Interestingly, we found that Tregs similarly accumulated as kidney tumor size increased (**Fig. 5A, bottom**), and that Treg abundance correlated well with macrophage-to-monocyte ratios in kidney as well as melanoma cancers (**Fig. 5B**). The positive correlation between Treg and macrophage density spurred us to ask whether one caused the other. Using *Foxp3*-DTR mice, we found that depletion of Tregs dramatically reduced the macrophage-to-monocyte ratio in mouse B16 tumors (**Fig. 5C, top**) as assessed by flow cytometry. This result was phenocopied by treatment of mice with an anti-CTLA-4 antibody that specifically depletes tumor Tregs^39^ (**Fig. 5C, bottom, S5B-C**).

To further examine how Tregs may be influencing monocyte and macrophage proportions, we performed scRNA-seq analysis on mouse tumor myeloid cells from B16 tumor-bearing control and *FoxP3*-DTR mice. *Csf1r*^+^ *Mafb*^+^ clusters from this experiment were aggregated with those from the original wild-type B16 tumor sample in **Figure 1** and we observed similar cell populations across both experiments and treatment conditions (**Fig. 5D, S5D-E**). Cluster proportions were modestly shifted with Treg loss (**Fig. 5D**), but cells from control and Treg- depleted tumors shared similar differentiation trajectories (**Fig. 5E**). However, Monocle analysis revealed differences in the accumulation of cells along the trajectory. Namely, while tumor monocytes, “Mono-Int”, and TAMs from the control sample acquired progressively increased pseudotime scores, “Mono-Int”, and TAM populations in the *Foxp3*-DTR sample did not exhibit sequential increases in pseudotime scores (**Fig. 5E**). In effect, TAM progression appeared stunted following depletion of Tregs.

Indeed, in addition to increased expression of inflammatory and immunomodulatory genes (e.g., *Ccl24*, *Arg1*, *Retnla*, *Mmp12*, *Mmp13, Nos2*), expression of monocyte-associated genes was sustained in TAMs from Treg-depleted tumors (**Fig. 5F-G, Table S3**). Moreover, expression of genes tied to macrophage differentiation (e.g., *C1qa*, *H2-Ab1*, *Apoe, Ms4a7*) were decreased across stages of differentiation (**Fig. 5H, S5F**), further indicating these TAMs were more immature. Our analysis suggests that the Treg-depletion may impair implementation of TAM transcriptional programs, a remodeling detected early during tumor monocyte differentiation. Altogether these findings support a model in which Treg abundance promotes an accumulation of terminally-differentiated TAMs in both mouse and human tumors.

### Multiparametric analysis of myeloid cell composition improves classification of kidney cancer patient antitumor responses

Given this association between T cell subset density and TAM maturation, we sought to further explore how features of tumor macrophage infiltration could be harnessed to reliably inform features of patient outcome, such as survival. Analysis of TCGA kidney cancer samples using a myeloid gene signature from CiberSort^64^ demonstrated that patients with varying levels of overall myeloid cell density did not significantly differ in their survival (**Fig. 6A, left**). We next stratified TCGA patients based on levels of monocyte/macrophage lineage genes *CSF1R* and *MAFB*, finding that patients with higher levels of these appreciated only modest improvements in outcome (**Fig. 6A, middle**). As these genes are not strictly macrophage-specific, we leveraged our scRNA-seq analyses of human kidney samples to generate signature scores based on the ratio between macrophage and monocyte (**Fig 4E**). However, no significant differences in survival were revealed using this metric (**Fig. 6A, right**).

**Figure 6.**
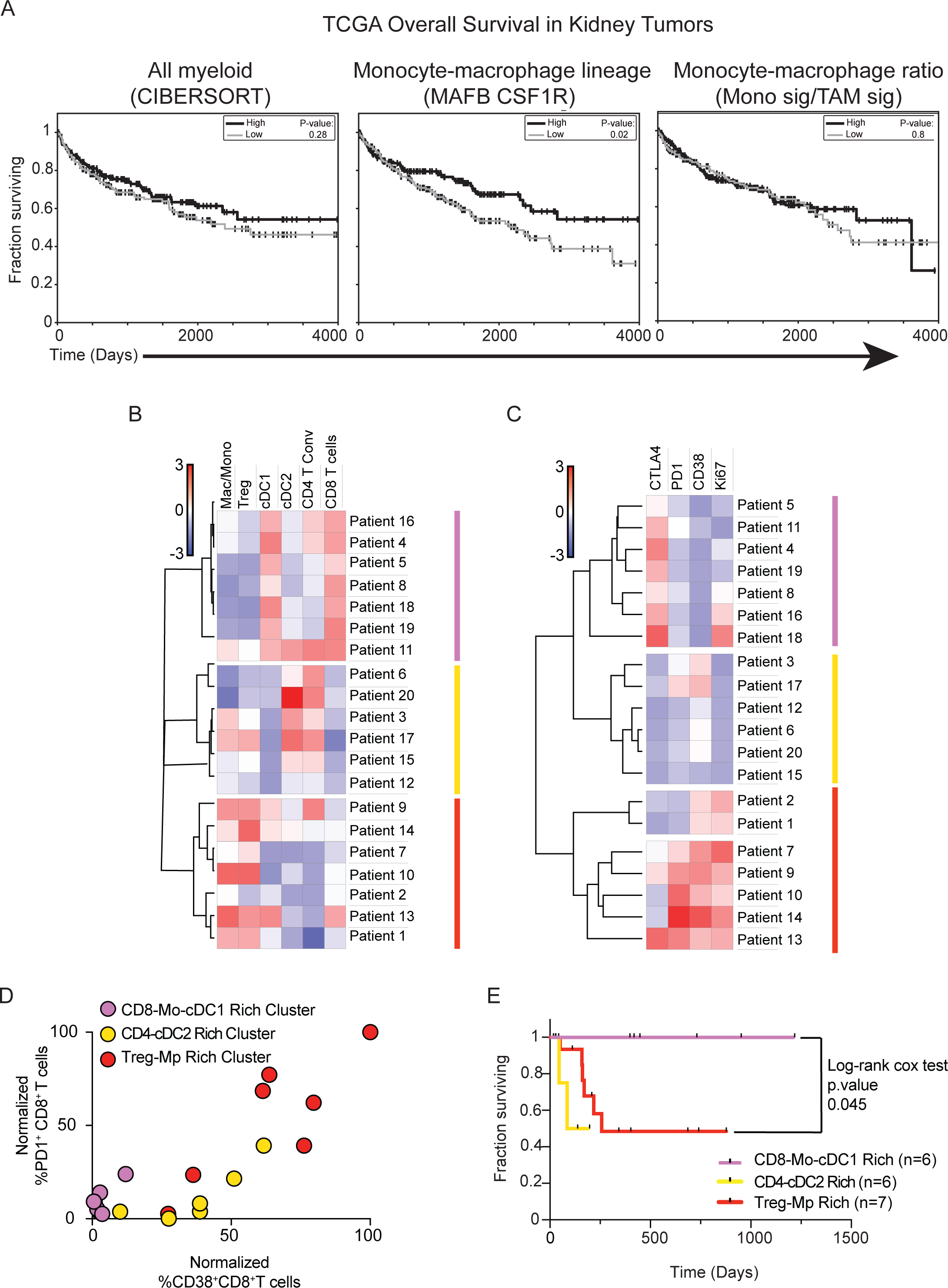
Multiparametric analysis of tumor myeloid composition identifies kidney cancer patients with effector CD8^+^ T cell responses and improved survival rates. **(A)** Survival curves of kidney tumor patients whose TCGA tumor samples exhibited high (33%) or low (33%) levels of expression levels of pan-myeloid gene signatures derived from CIBERSORT (**left**), *MAFB* and *CSF1R* (**middle**), or ratio of monocyte-to-TAM gene signatures (Fig. 4) (**right**). **(B)** Heatmap of specified immune cell population frequencies detected in human kidney tumor samples by flow cytometry. **(C)** Heatmap of specified surface receptor or Ki-67 expression frequencies amongst CD8^+^ T cells from human kidney tumor samples that were analyzed with flow cytometry. **(D)** Quantification of the frequency of CD8^+^ T cells from human tumor kidney samples that are PD1^+^ or CD38^+^. Labeling of dots corresponds to patient groups (**B,C**). **(E)** Survival curves of kidney cancer patients in cohort analyzed (**B-D**).

As TAM density did not appear to robustly inform patient outcome, we sought to test how TAM abundance corresponds with other immune parameters and may stratify patients. We thus analyzed a cohort of kidney cancer patient biopsies by flow cytometry and quantified immune cell population frequencies. Unbiased clustering analysis of samples revealed three groups of patients that exhibited distinct immune composition patterns (**Fig. 6B**). As suggested from our previous analysis (**Fig. 5A**) macrophage/monocyte ratio and Treg density represented a strong classifier which delineated the patients and revealed a group that was highly enriched for both macrophages and Tregs (Treg-Mp; **red**). Although the other groups were generally TAM/monocyte^lo/int^, one group (CD8-Mo-cDC1; **pink**), but not the other (CD4-cDC2; **yellow**), was distinguished by notable infiltration of cDC1, which are critical for CD8^+^ T cell responses ^37, 38, 40^, and that group presented with uniformly high CD8^+^ T cell infiltration. Further, the same patients clustered together based on expression levels of checkpoint regulators (i.e., PD-1, CTLA-4, CD38) and proliferative capacity (i.e. Ki-67) within the CD8^+^ T cell compartment (**Fig. 6C**). Notably, CD8^+^ T cells from the CD8-Mo-cDC1 group (**pink**) expressed low levels of exhaustion markers PD-1 and CD38 (**Fig. 6C-D**) and were also distinguished by higher CTLA-4 protein expression, which may indicate ongoing activation^65^ (**Fig. 6C-D**). In contrast, the Treg- Mp (**red**) group showed the highest levels of both exhaustion markers and Ki-67.

In testament to the heightened antitumor CD8^+^ T cell profile associated with low macrophage and Treg abundance but high cDC1 density, the subset of patients with these attributes (**pink**) appreciated dramatically improved survival (**Fig. 6E**). Moreover, we found that measuring the ratio of cDC1s to macrophages through combined gene signatures rather than use of either signature alone (**Fig. S6A-C)** allowed for identification of kidney cancer patients with better survival in an external bulk RNAseq dataset (TCGA). Thus, fine-tuned stratification of kidney cancer TIME provided the resolution critical for identifying distinct patient classes including this CD8-Mo-cDC1 group, which defines patient with the best anti-tumor immune response regardless of tumor stage and treatment.

## Discussion

Here we undertook scRNA-seq analysis of tumor monocytes and macrophages to determine the key hallmarks of their transcriptional diversity. We found two types of differentiation to manifest during tumor development. On the one hand we found a classical lineage differentiation trajectory that progresses from monocytes-to-macrophages in a way that has been long appreciated^66^ with a discernable ‘intermediate’ monocyte (“Mono-Int”) cell population. A “Mono-Int” population is, for reference, well-described in other settings. For example, Randolph and colleagues detect ‘intermediate’ monocytes in lymphoid and non-lymphoid tissue in steady-state conditions^67^, and fluorescent real-time lineage tracing identifies cells undergoing that transition during allergic challenge^68^.

On the other hand, we found two differentiation layers – ‘stress-responsive’ and ‘IFN-responsive’ – that co-exist along that trajectory and that were shared across multiple mouse models as well as a profiled human kidney cancer biopsy (**Fig. 1, 4**). These programs were also present in other recently published studies^18, 60, 69^. For example, in a pan-cancer study, Cheng et al. discern myeloid populations whose primary distinction is their expression of IFN-induced genes (e.g., *ISG15*^+^ TAMs). A notable difference in our interpretation compared to these reports lies in our incorporation of these layers within the monocyte-macrophage differentiation axis, rather than proposing them as a unique trajectory. Through independent profiling of purified monocytes and macrophages in our study and pseudotime analysis (**Fig. 1-2**), we find the stress-responsive signatures evident in both cell populations and indeed across them. In additional support of such a view, we found that an IFN-responsive signature was enriched amongst monocytes in one mouse model and macrophages in another (**Fig. 1, 3**). We believe that this represents that macrophages can differentiate in two dimensions – progression through the classical lineage as well as acquisition of specialized states characterized by examples of IFN or stress exposure. Intuitively, this is similar to CD4^+^ T cells that can differentiate along a naïve-effector-memory axis while also being able to layer on Th1/Th2/Th17 programs^70^.

Despite the latter comparison, we notably do not find any populations, nor indeed any cells, that have an exclusively “M1” or “M2” signature (**Fig. 3**). Individual genes such as *Arg1* are associated with certain clusters, as some have observed^18^, but both correlation and signature analyses fail to identify any of the described ‘M1’ or ‘M2’ genes as either being selectively linked with one another in single cells, or as key classifiers of cell clusters. To this extent, the ‘M1/M2’ nomenclature has provided direction in the fruitful study of myeloid cell signaling and differentiation but does not appear to be accurately categorize distinct differentiation states, at least for tumors *in vivo*. We note the absence of data to the contrary of this conclusion in other recent reports^55, 58, 69^ although of course individual nomenclature (e.g., “M2-like”) is clearly a matter of choice and needs only discussion as to which part of the *in vitro* signature might be biologically relevant.

One important aspect of myeloid biology that requires further elaboration is how to identify IFN- and stress-responsive phenotypes. For example, Gubin et al. use iNOS as a marker by flow cytometry to define the IFN-stimulated population induced by ICB whereas Cheng et al. utilize *ISG15*. Particularly in the former study which studied macrophage identity post checkpoint blockade therapies, varied levels of type I and II IFNs may also modulate properties of this differentiation layer. In the case of ‘stress-responsive’ populations, our data also point to IL-7R*α* expression, which may indicate involvement of TSLP signaling through heterodimeric pairing with TSLPR^71, 72^. In human macrophages, SEPP1 was present in our study as well as in Cheng et al., but this or its homolog is not formative for what appears to be the analogous population in mice. An important set of conserved genes for ‘stress-responsive’ macrophages, taken from our manuscript, is their consistent and significant enrichment for glycolytic genes, particularly in comparison to conventional C1q ‘mature’ TAMs. Given that HIF- 1 is known to induce glycolytic genes in hypoxic conditions^73^, this finding raises the questions of whether these cells are selected for hypoxic environments where oxidative phosphorylation may not proceed, as well as their specific function. Going forward, the use of multiplexed imaging technologies such ion beam imaging (MIBI) ^74, 75^ and single-cell spatial transcriptomics^76–78^ will enable this analysis.

Our investigation of monocyte/macrophage differentiation led us to explore how its regulation could inform our understanding of antitumor immunity. Analysis of kidney cancer and melanoma patient cohorts revealed an increase in macrophage-to-monocyte ratios with tumor grade, a rise that coincided with Treg density and was Treg-dependent. Tregs exert potent immunosuppression and are thought to restrain T cell activity and antitumor responses through modulation of DC stimulatory capacity, production of immunosuppressive cytokines and substrates, and competitive usage of growth factors and metabolic byproducts^39, 79–81^. It is becoming clear now that tumor Tregs also strongly influence the monocyte/macrophage lineage, likely through multiple mechanisms. In a recent study, tumor Tregs promoted tumor macrophage numbers by supporting their mitochondrial capacity and viability^82^. Here our scRNA-seq data demonstrates that early-stage monocytes and “Mono-Int” cells are already unable to properly implement TAM-associated transcriptional programs in the absence of Tregs, indicating that Tregs also fuel macrophage differentiation processes. This liaison between Tregs and macrophages mirrors one identified elsewhere such as adipose fat of lean mice where Tregs are thought to actively maintain homeostasis and hold inflammatory macrophages at bay^83, 84^. Similarly, during the resolution of injury and inflammation in skeletal muscle and heart tissue, a transition from pro- to anti-inflammatory macrophages occurs in a manner that appears to rely on Treg accumulation^85–88^. That Tregs may act on tumor macrophages in a similar fashion offers another example of how the TIME can exploit immune programs of “accommodation” that are otherwise in place to achieve tissue homeostasis in the face of perturbations^89^.

Accumulation of a broad swath macrophages in the TIME has previously been implicated with poor outcome^90^. Consistent with this but at higher resolution, we detected a group of kidney cancer patients for whom high macrophage-to-monocyte abundance was associated with diminished T cell infiltration and exhaustion of those cells detected, concurring with other very recent reports^58, 60^. Our manuscript thus points to an emerging trio of Tregs, macrophages, and exhausted T cells, whereby effector T cells may be corrupted through direct cellular interactions with TAMs, as has been suggested by observations of TAM-CD8^+^ T cell co- localization in ccRCC^60^, or indirectly through macrophage-induced Treg expansion and activity^15, 91^ or DC suppression^39, 92^.

Yet, high myeloid infiltration or skewed macrophage-to-monocyte ratios alone were not prognostic for KIRC patient survival. Indeed, although macrophages have often been found to be negatively associated with patient outcome, macrophage abundance as a sole biomarker has not been universally useful with prior studies similarly reporting instances in which macrophage abundance is not informative for patient cohorts with specific cancer sub-types, treatment regimens, or tumor stage^93–96^. Clustering analysis of kidney TIME composition using comprehensive immune parameters, however, uncovered an archetype characterized by low macrophage-to-monocyte in conjunction with high cDC1 infiltration. These patients (CD8-Mo- cDC1) had elevated infiltration of CD8^+^ T cells with low surface expression of proteins associated with exhaustion and appreciated highly enhanced survival rates (**Fig. 6, pink**).

Notably, recent work focused on ascertaining the different immune archetypes across solid tumors suggests that these patient groups span cancer type^51^.

Identification of a CD8-Mo-cDC1 archetype emphasizes the value of integrating multiparametric biomarkers as a means to better parse patient outcome and to establish principles of TIME organization. Given that T cell activity appears to be collectively influenced by multiple immune cell populations with distinct partnering patterns, our analysis suggests that dual targeting of TIME axes may elicit the best CD8+ T cell responses. For example, reprogramming and/or depletion of macrophages may relieve active suppression^5, 44^ and strategies that boost cDC1 recruitment and survival^2^ may further benefit even those with favorable macrophage-to-monocyte density. It is also notable that this protective archetype is specifically enriched for monocytes. Indeed, monocyte differentiation into macrophages may not be inevitable and accumulation of “Mono-Int” cells have been detected in multiple forms of inflammation^17, 29, 97–100^. Additionally, the potential importance of monocytes is indicated by their increased numbers in the blood of ICB responsive as compared to non-responsive melanoma patients^50^. In ccRCC patients, IFN-responsive TAMs exhibited lower levels of *HLA-DR*, reminiscent of “Mono-Int” cells described here, and higher levels of these TAM-ISG^hi^ were predictive of survival after TKI treatment^94^. Such a relationship opens questions across cancer type; namely, whether “Mono-Int” are distinct in their antitumor function, and how might monocytes be additive or synergistic with cDC1 to drive antitumor CD8 T cells?

Altogether these findings contribute to the endeavor of clarifying useful distinctions in myeloid gene expression and highlights settings in which multiparametric analysis of tumor myeloid composition can inform patient immune archetype and guide development of relevant therapies.

## Methods

### Mice

The following mice were housed and/or bred under specific pathogen-free conditions at the University of California, San Francisco Animal Barrier Facility: C57BL/6J (The Jackson Laboratory), MMTV-PyMT-mCherry-OVA transgenic^101^, and *Foxp3*-DTR (The Jackson Laboratory). All mice were handled in accordance with NIH and American Association of Laboratory Animal Care standards, and experiments were approved by the Institutional Animal Care and Use Committee of the University of California, San Francisco.

### Human Tumor Samples

RCC or melanoma tumor samples were transported from various cancer operating rooms or outpatient clinics. All patients consented by the UCSF IPI clinical coordinator group for tissue collection under a UCSF IRB approved protocol (UCSF IRB# 20-31740). Samples were obtained after surgical excision with biopsies taken by Pathology Assistants to confirm the presence of tumor cells. Patients were selected without regard to prior treatment. Freshly resected samples were placed in ice-cold DPBS or Leibovitz’s L-15 medium in a 50 mL conical tube and immediately transported to the laboratory for sample labeling and processing. The whole tissue underwent digestion and processing to generate a single-cell suspension. In the event that part of the tissue was sliced and preserved for imaging analysis, the remaining portion of the tissue sample was used for flow cytometry analysis.

### Tumor Cell Lines

B16-F10 cells (ATCC, CRL-6475) were purchased and cultured at 37°C in 5% CO2 in DMEM (Invitrogen), 10% FCS (Benchmark), Penicillin, Streptomycin, and L-Glutamine (Invitrogen). B16-F10-ZsGreen was previously generated in our laboratory as described^102^.

### Mouse Tumor Cell Injections and Growth

Prior to injection, adherent B16-F10 or B16-ZsGreen tumor cells were dissociated with 0.05% Trypsin-EDTA (Thermo Fisher Scientific) and washed 2-3X with DPBS (Thermo Fisher Scientific). 1.0x10^5^ – 2.5x10^5^ cells were resuspended in DPBS and mixed 1:1 with Matrigel GFR (Corning). Mice were injected subcutaneously with a volume of 50 μl either unilaterally or bilaterally depending on the experimental setup. Tumor tissue was harvested approximately 12- 16 days later.

MMTV-PyMT-mCherry-OVA transgenic mice were bred and genotyped for the transgene by PCR. Spontaneous tumor growth was monitored, and tumors were harvested when the mice were approximately 20-30 weeks of age.

### Mouse Tissue Processing and Flow Cytometry Staining

Mouse tumor tissue was harvested and enzymatically digested with 0.2mg/ml DNase I (Sigma-Aldrich), 100U/ml Collagenase I (Worthington Biochemical), and 500U/ml Collagenase Type IV (Worthington Biochemical) for 30 minutes at 37°C while under constant agitation. Blood was collected via cardiac puncture from mice that were euthanized by overdose with 2.5% Avertin. Blood samples were treated with 175 mM NH4Cl for 5 minutes at room temperature to lyse red blood cells.

Samples were filtered, washed with stain media (DPBS, 2% FCS), and resuspended again with stain media. Cells from this single cell suspension were washed with DPBS and stained with Zombie NIR fixable viability dye (BioLegend) for 30 minutes at 4°C. Cells were washed and resuspended with stain media containing anti-CD16/32 (BioXCell), 2% rat serum, 2% Armenian hamster serum, and antibodies against surface proteins of interest. Cells were stained for 30 minutes at 4°C. At times cells were then washed and stained for intracellular protein levels, for which they were fixed, permeabilized, and stained according to BD Cytofix/Cytoperm Kit (BD Biosciences) or the FoxP3/Transcription Factor Staining Buffer Set (Thermo Fisher).

The following antibodies were from Biolegend: anti-mouse CD45, anti-mouse Ly-6C (HK1.4), anti-mouse CD11b (M1/70), anti-mouse CD11c, anti-mouse MHC-II, anti-mouse F4/80, anti-mouse CD24, anti-mouse Ly-6G, anti-mouse NK1.1, anti-mouse CD90.2, anti-mouse/human CD45R/B220, anti-mouse CD301b, anti-mouse CD64, anti-mouse CD127. The following antibodies were from BD Biosciences: anti-mouse Siglec-F, anti-mouse CD106. The following antibodies were from R&D: anti-mouse/human ARG1, normal sheep IgG. The following antibodies were from ThermoFisher: anti-mouse FoxP3.

Following staining, cells were washed, resuspended in stain media, and analyzed on a BD Biosciences Fortessa or sorted with a BD Biosciences FACSAria Fusion. Flow cytometry data was analyzed using FlowJo software (BD Biosciences).

### Human Tissue Processing and Flow Cytometry Staining

Tumor or metastatic tissue was thoroughly chopped with surgical scissors and transferred to gentleMACS C Tubes (Miltenyi Biotec) containing 20 uL/mL Liberase TL (5 mg/ml, Roche) and 50 U/ml DNAse I (Roche) in RPMI 1640 (Invitrogen) per 0.3 g tissue. gentleMACS C Tubes were installed onto the gentleMACS Octo Dissociator (Miltenyi Biotec) and incubated for 45 minutes according to the manufacturer’s instructions. Samples were then quenched with 15 mL of sort buffer (DPBS, 2% FCS, 2mM EDTA), filtered through 100 μm filters, and spun down. Red blood cell lysis was performed with 175 mM ammonium chloride if needed.

Cells were incubated with Human FcX (Biolegend) to prevent non-specific antibody binding. Cells were then washed in DPBS and incubated with Zombie Aqua Fixable Viability Dye (Thermo). Following viability dye, cells were washed with sort buffer and incubated with cell surface antibodies that had been diluted in the BV stain buffer (BD Biosciences) for 30 minutes on ice. Cells were subsequently fixed in either Fixation Buffer (BD Biosciences) or in Foxp3/Transcription Factor Staining Buffer Set (eBioscience) if intracellular staining was required. The following antibodies were from BD Biosciences: anti-human HLA-DR, anti-human CD56, anti-human CD127, anti-human CD25, anti-human CD45RO, anti-human PD-1, anti-human CTLA-4, and anti-human CD64. The following antibodies were from ThermoFisher: anti-human CD45, anti-human CD3*ε*, anti-human FoxP3, anti-human Ki-67, anti-human CD19, anti-human CD20, anti-human CD56, and anti-human CD11c. The following antibodies were from Biolegend: anti-human CD4, anti-human CD8*α*, anti-human CD38, anti-human CD16, CD1C/BDCA-1, anti-human CD14, anti-human CD304, and streptavidin. Anti-human BDCA-3 was purchased from Miltenyi,

Stained cells were washed and analyzed on a BD Biosciences Fortessa or sorted with a BD Biosciences FACSAria Fusion. Flow cytometry data was analyzed using FlowJo software (BD Biosciences).

### Single Cell RNA-Sequencing Data Generation

Sorted cells were resuspended at a concentration of 1x10^3^ cells/μl in media (DPBS, 0.04% BSA) and loaded onto the Chromium Controller (10X Genomics). Samples underwent single-cell encapsulation and cDNA library preparation using the Chromium Single Cell 3’ v1 or v2 Reagent Kits (10X Genomics). The cDNA library was sequenced on an Ilumina HiSeq 4000 (Illumina).

### Single Cell RNA-Sequencing Data Processing

Sequencing data was processed using 10X Genomics Cell Ranger V1.2 pipeline. Fastq files were generated from Ilumina bcl files with the Cell Ranger subroutine *mkfastq*. Fastq files were then processed with Cell Ranger’s *count* to align RNA reads against UCSC mm10 or GRCh38 genomics for mouse and human cells, respectively, using the aligner STAR^103^. Redundant unique molecular identifiers (UMI) were filtered, and a gene-cell barcode matrix was generated with *count*. *Mkfastq* and *count* were run with default parameters.

For mouse B16 tumor samples, the gene-cell barcode matrix was passed to the R software package Seurat (v2.3.0)^104^ for all downstream analyses. Genes that were expressed in at least 3 cells were included. Cells that did not express at least 200 genes were excluded, as were those that contained >5% reads associated with cell cycle genes^105, 106^. For mouse PyMT and human RCC tumor samples, raw feature-barcode matrices were loaded into Seurat (v3.1.5)^107^ and genes with fewer than 3 UMIs were dropped from the analyses. Matrices were further filtered to remove events with greater than 20% percent mitochondrial content, events with greater than 50% ribosomal content, or events with fewer than 200 total genes. The cell cycle state of each cell was assessed using a published set of genes associated with various stages of human mitosis^108^.

Using Seurat’s *ScaleData* function, read counts were log2 transformed and scaled using each cell’s proportion of cell cycle genes as a nuisance factor. A set of highly variable genes was generated by Seurat’s *FindVariableGenes* function, which were used for principal component (PC) analysis. Genes associated with PCs (selected following visualization with scree plots) were used for graph-based cluster identification and dimensionality reduction using *t*-distributed stochastic neighbor embedding (*t*-SNE) analysis. Seurat’s *FindAllMarkers* function was used for subsequent cluster-based analyses, including cluster marker identification and DE gene analyses.

### Single Cell RNA-Sequencing Signature Generation

To generate mouse monocyte- and macrophage-specific gene signatures (**Fig. 1**), sorted monocyte, TAM1, and TAM2 samples were aggregated, log2 transformed, and scaled using Seurat. DE gene analysis was performed using *FindMarkers* with the parameters log *N* fold change > 1.5 and a min.pct of 0.25. Genes were selected by ranked fold change and the criteria that min.pct1/min.pct2 > 1.5. Genes used for cell cycle regression analysis were excluded. The top 10 genes (or fewer if less remained) were median normalized and aggregated, scaled 0-1, and plotted on specific *t*-SNE plots.

Gene signatures for cellular programs such as metabolism^57^, “M1” and “M2” polarization^55^, and MHC-II-associated genes (GSEA, REACTOME_MHC_CLASS_II_ANTIGEN_PRESENTATION), previously published cell types^53, 59^, or cell populations identified here were generated by taking the mean of log-normalized expression for a particular set of genes related to the specific pathway or phenotype. To visualize the distribution of these scores across cells, we binarized the distribution of the score at the 70th percentile unless specified otherwise and overlaid on the calculated *t*-SNE coordinates.

For correlation analysis of “M1” and “M2” genes, the expression of each gene in the signatures was calculated for each B16 tumor *Csf1r*^+^ *Mafb*^+^ cluster cell and binarized at the median. A cross-correlation gene-gene matrix was obtained using the R *cor* function with *method=”pearson”*.

### Single Cell RNA-Sequencing Sample Aggregation

To perform pairwise comparison analyses between B16 tumor myeloid cell clusters from wildtype and Treg-depleted mice, the objects were first transformed from Seurat v2 to Seurat v3. The raw UMI counts were renormalized using person residuals from “regularized negative binomial regression,” with sequencing depth a covariate in a generalized linear model via the R sctransform package^109^. Pairwise “anchor” cells were identified between the three objects using the wild-type mouse from **Figure 1** as a reference via the Seurat *FindIntegrationAnchors* function. Briefly, each pair of samples was reduced to a lower dimensional space using diagonalized Canonical Correlation Analysis (CCA) using the top 3000 genes with the highest dispersions. The canonical correlation vectors were normalized using L2-normalization. Multiple Nearest Neighbors (MNNs) across datasets were identified for each cell in each dataset and cell-cell similarities are used as anchors to integrate the datasets together using the Seurat *IntegrateData* function.

### Single Cell RNA-Sequencing Pseudotime Analysis

Raw UMI counts from the cleaned and processed Seurat objects for the control and Treg- depleted mouse experiment were extracted and coerced into Monocle2^54, 110^. *CellDataSet* objects, normalizing the data using a negative binomial distribution with fixed variance (*negbinom.size*). Each object was independently processed to identify a pseudotime trajectory. Briefly, each object was processed to estimate per-cell coverage and sequencing depth (*estimateSizeFactors*) and gene dispersions (*estimateDispersions*). Cells were further filtered to retain high-quality cells with >=500 genes and genes were filtered to retain only those in at least 10 cells. The dataset was reduced to 2 dimensions using the *DDRTree* algorithm and the marker genes that differentiated the *Ly6c2*^+^ *Hp*^+^ monocytes and *C1qa*^+^ TAM clusters from other cell types were used to guide the trajectory inference. Relative pseudotime was obtained through a linear transformation relative to the cells with the lowest and highest pseudotimes *(1- min_pseudotime)/max_pseudotime*. The “wave” plots in **Figure 1** were constructed using the Seurat LogNormalized counts and the relative pseudotime described above for the top 5 genes per cluster as identified by scRNA-seq.

### TCGA Survival Analyses

Tumor RNAseq counts and transcripts-per-million (TPM) values for kidney renal clear cell carcinoma (KIRC) samples from the Toil recompute data in the TCGA Pan-Cancer (PANCAN) cohort^111^ were downloaded from the UCSC Xena browser^112^. A gene signature score for each patient was calculated using the gene signature score method below. The feature gene signature scores were calculated using an m x n matrix where m represented the TPM normalized log2 counts per million (logCPM) expression of the feature signature genes and n represented the selected sample set^113^. The expression of each gene was converted to percentile ranks across the samples using the SciPy Python module^114^. The top and bottom 30 percentile were then used to define low and high patients for each respective signature unless otherwise noted.

### *In Vivo* Mouse Treatments

To deplete Treg cells, *Foxp3*-DTR and control mice were injected intraperitoneally with 500ng of unnicked diptheria toxin (List Biologics, 150). Mice were typically injected 9, 10, and 12 days following initial inoculation with tumor cells.

For specified experiments, wild-type mice were injected intraperitoneally 7, 9, 10, 11, and 13 days following tumor injection with 250 µg of anti-mouse CTLA-4 IgG2c (modified clone 9D9, Bristol-Myers-Squibb), mouse IgG2C isotype, anti-mouse CTLA-4 IgG1 (modified clone 9D9, Bristol-Myers-Squibb), or mouse IgG1 isotype.

### Statistical analysis and data visualization

Comparisons between groups were analyzed using GraphPad Prism software. Experimental group allocation was determined by genotype or by random designation when all wild-type mice were used. Error bars represent mean ± SEM calculated with Prism unless otherwise noted.

Comparisons between 2 groups were analyzed with Student’s *t*-test. For statistical measures between more than 2 groups, one-way ANOVA was performed unless otherwise noted.

Nonsignificant comparisons are not shown. Investigators were not blinded to experiment group assignment during experimental data generation or analyses. The R packages Seurat and ggplot2 were used to generate figures.

## Acknowledgments

We would like to thank E. Wan and the Institute for Human Genetics at UCSF for helping prepare samples for scRNA-seq as well as the UCSF Flow Cytometry Core for maintenance of flow cytometers and sorters. We would like to also thank J.J. Engelhardt and Bristol-Myers-Squibb for Fc-modified anti-CTLA-4 antibodies. This work was supported by the US National Institutes of Health (R01CA197363 and U01CA217864 to M.F.K). A.M.M. is supported by the Cancer Research Institute as a Cancer Research Institute/Amgen Fellow. Acquisition and analysis of certain human samples was partially funded by contributions from Abbvie, Amgen, Bristol-Myers Squibb, Genentech, and Pfizer.

## Author Contributions

A.M.M, A.J.C., M.B., and M.F.K. designed experiments. A.M.M, A.J.C., and M.B. performed experiments unless specified. A.R.R. and J.L.P. participated in the processing and analysis of scRNA-seq. B.S. participated in the analysis of TCGA samples. J.T., A.B., R.J.A., M.K.R, and K.C.B. participated in experimental data collection and flow cytometry analysis. V.C. managed the acquisition of human tumors samples. A.M.M., A.J.C., and M.F.K. wrote the manuscript. A.M.M., A.J.C., M.B., A.B., and R.J.A. edited the manuscript.

## Declaration of Interests

M.F.K. is a founder and shareholder in Pionyr Immunotherapeutics that develops novel immunotherapeutics that target and tune myeloid cells. M.B and J.L.P. are shareholders in Pionyr Immunotherapeutics.

---

**Supplementary Figure 1.**
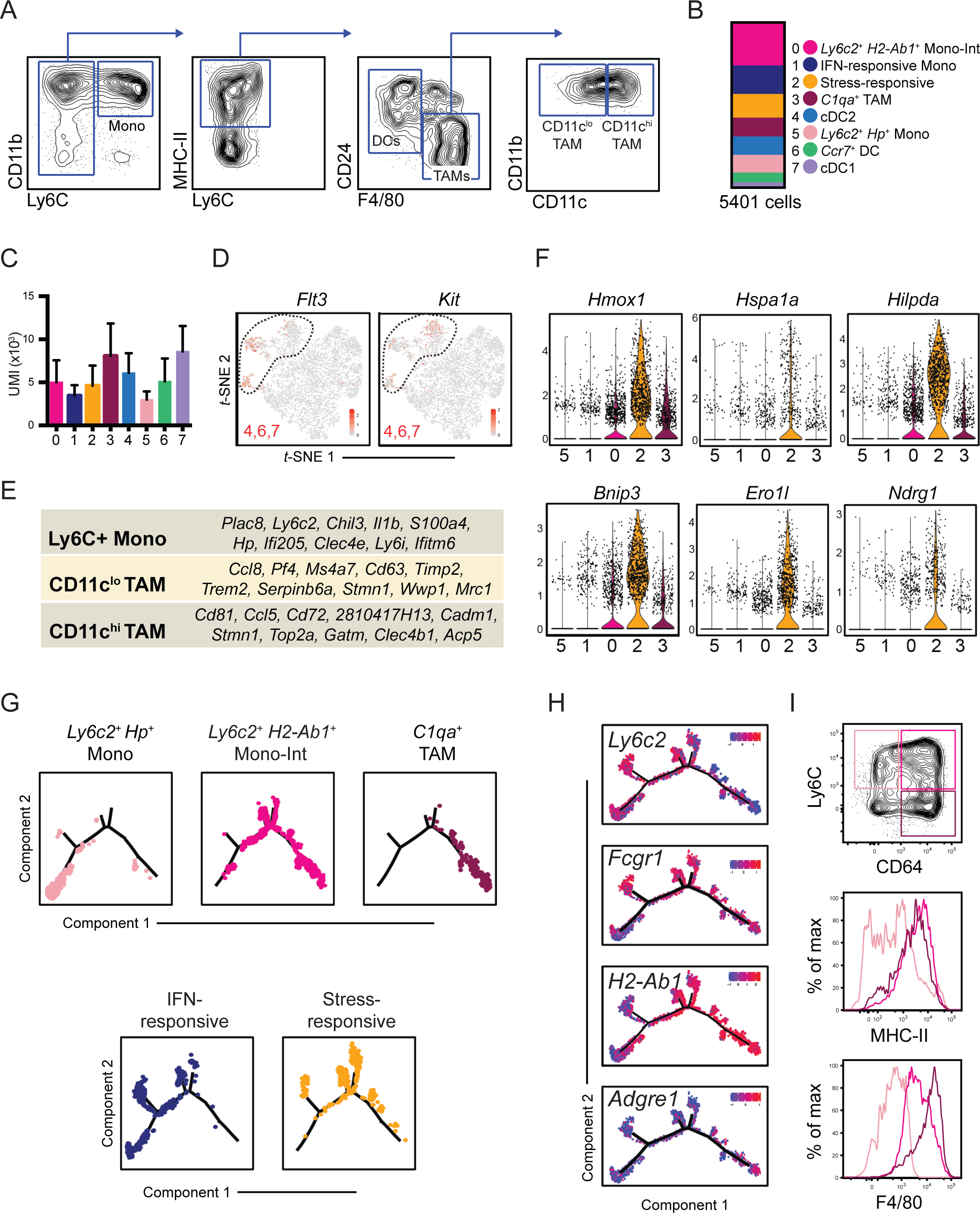
(A) Flow cytometry gating schematic for sorting and analysis of specified B16 tumor myeloid populations. **(B)** Quantification of cells recovered from scRNA-seq analysis of B16 bulk tumor myeloid cells (Fig. 1B). **(C)** Quantification of unique molecular identified (UMI) counts in each myeloid cell cluster (Fig. 1B). **(D)** Expression of *Flt3* (**left**) and *Kit* (**right**) displayed on t-SNE plot of B16 tumor myeloid cells (Fig. 1B). **(E)** List of cell type-specific DE genes identified from comparative analysis of Ly6C^+^ monocyte, CD11c^lo^ TAM1, or CD11c^hi^ TAM2 populations that were individually sorted and processed for scRNA-seq analysis. (Fig. 1A). **(F)** Expression levels of selected stress-response genes amongst *Csf1r*^+^ *Mafb*^+^ cell clusters (Fig. 1C). **(G)** Differentiation trajectory model of *Csf1r*^+^ *Mafb*^+^ cells (Fig. 1G) with display of cells from individual clusters (Fig. 1C). **(H)** Expression of selected canonical monocyte- and macrophage-related genes displayed on the differentiation trajectory plot (Fig. 1G). Flow cytometric gating of B16 tumor-infiltrating monocytes (pink; Ly6C^+^ CD64^-^), “Mono- Int” (magenta; Ly6C^+^ CD64^+^), and macrophages (purple; Ly6C^-^ CD64^+^) (**top**), and their surface protein expression of MHC-II (**middle**) and F4/80 (**bottom**),

**Supplementary Figure 2.**
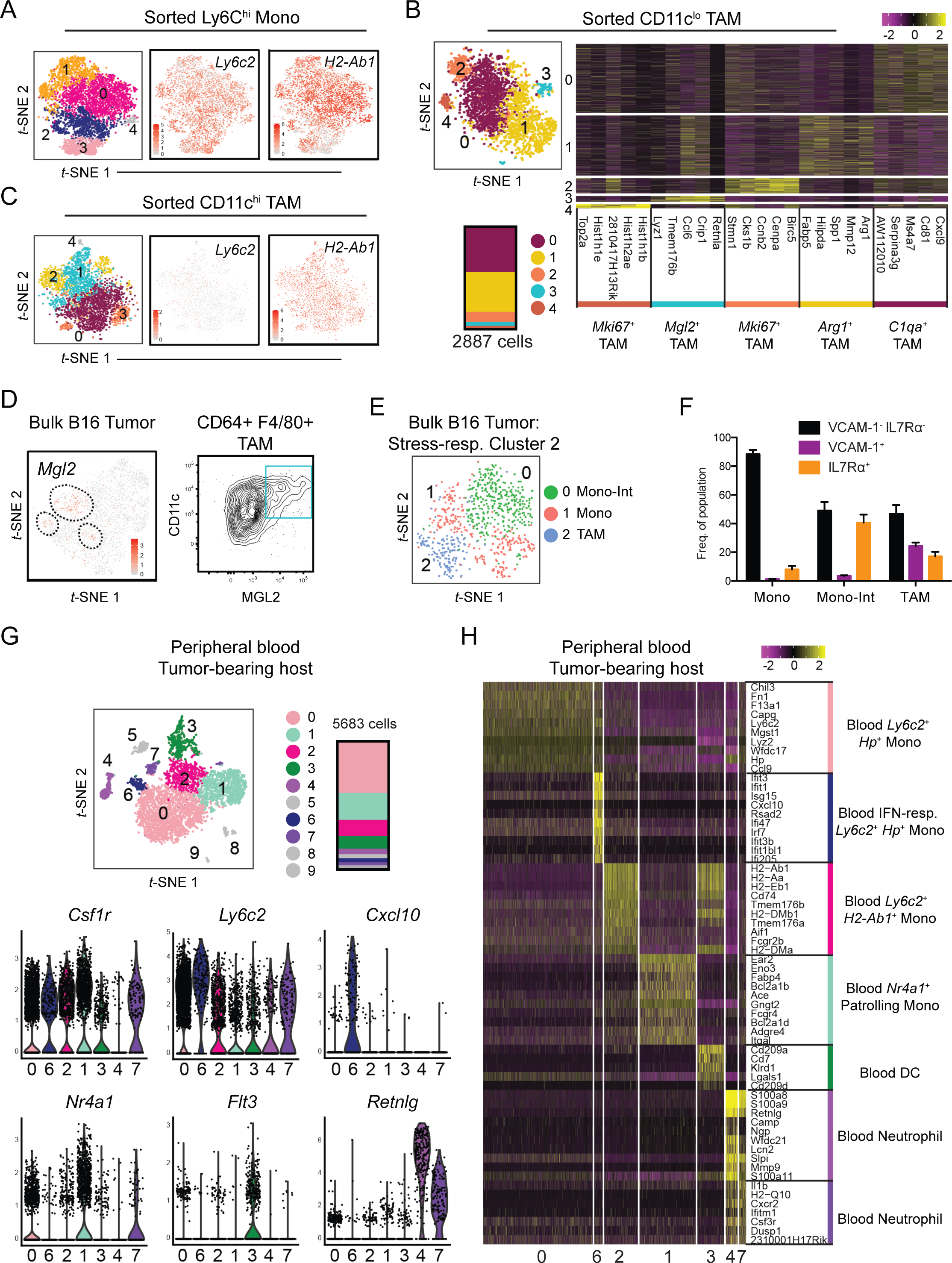
(A) *t-SNE* plot of sorted Ly6C^+^ monocytes sorted from B16 tumors (**left;** Fig. 2A) with expression of *Ly6c2* (**middle**) and *H2-Ab1* (**right**) displayed. **(B)** *t*-SNE plot of graph-based clustering (**left, top**) and quantification (**left, bottom**) of CD11c^lo^ TAMs sorted from B16 tumors and processed for scRNA-seq, and heatmap showing expression levels of top 5 DE genes between clusters **(right**). Genes are ranked by fold change. **(C)** *t-SNE* plot of sorted CD11c^hi^ TAMs sorted from B16 tumors (**left;** Fig. 2B) with expression of *Ly6c2* (**middle**) and *H2-Ab1* (**right**) displayed. **(D)** Expression levels of *Mgl2* displayed on t-SNE plot of bulk *Csf1r*^+^ *Mafb*^+^ cells sorted from B16 tumors (**left;** Fig. 1C), and MGL2 surface protein levels on TAMs from B16 tumors (**right**). **(E)** *t-SNE* plot of graph-based clustering analysis of stress-responsive cluster (Cluster 2) from bulk myeloid cells (Fig. 2C). **(F)** Quantification of frequency of VCAM-1^-^ IL7R*α*^-^, VCAM-1^+^, or IL7Ra^+^ cells detected within B16 tumor monocyte, Mono-Int, or TAMs by flow cytometry. Data is pooled from 2 independent experiments, each with 3-5 mice (mean ± SEM). **(G)** *t*-SNE plot of graph-based clustering of myeloid cells sorted from peripheral blood of B16 tumor-bearing mice and processed for scRNA-seq (**top**), and expression levels of selected genes amongst myeloid clusters (**bottom**). **(H)** Heatmap displaying expression levels of top 10 DE genes between blood myeloid cell clusters (**G**) with genes ranked by fold change.

**Supplementary Figure 3.**
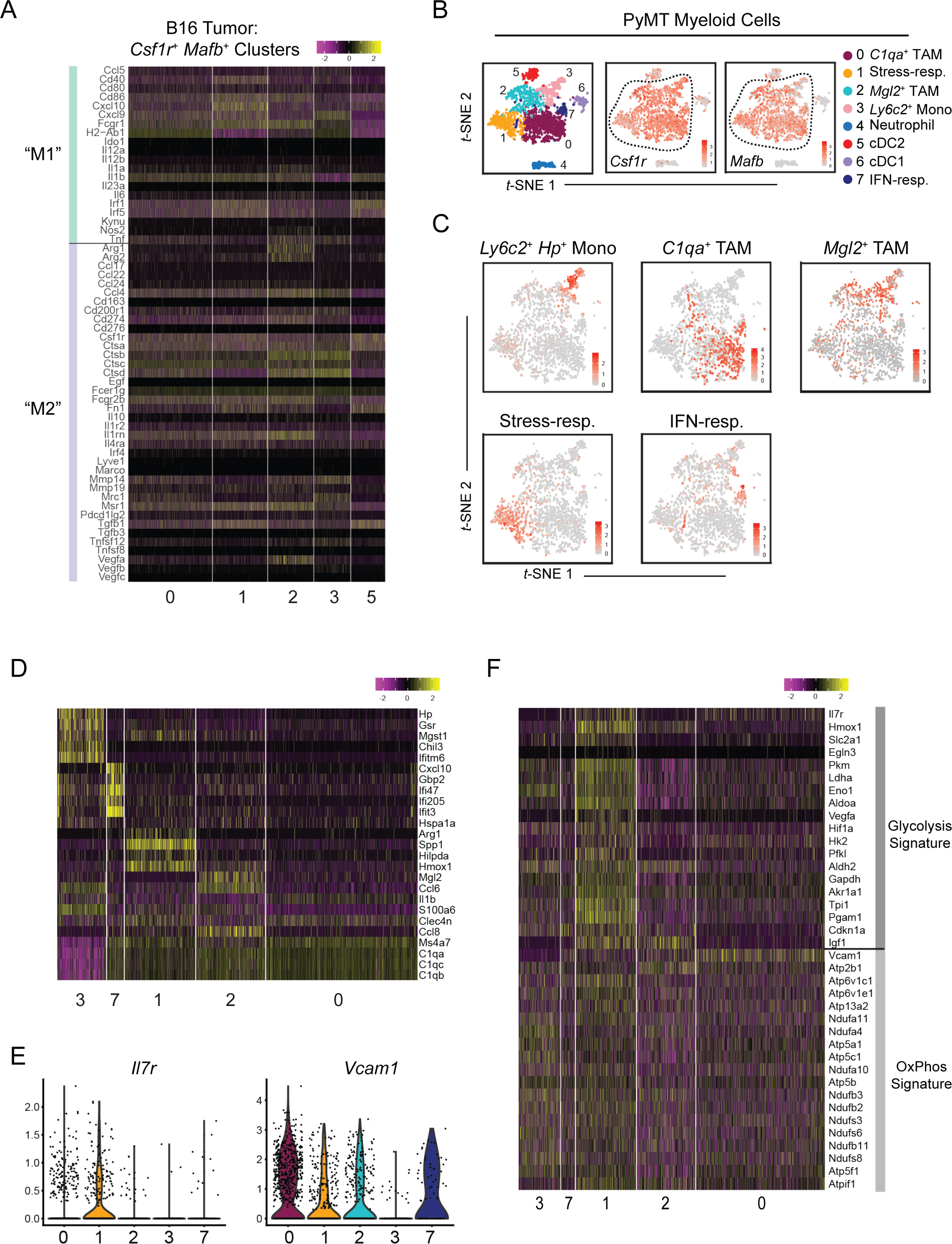
(A) Heatmap displaying expression levels of selected “M1”- and “M2”-associated genes by *Csf1r*^+^ *Mafb*^+^ cluster cells from B16 tumors (Fig 1C). **(B)** *t*-SNE plot and graph-based clustering of myeloid cells sorted from PyMT tumors and processed for scRNA-seq (**left**) with expression levels of *Csf1r* (**middle**) and *Mafb* (**right**) displayed. **(C)** Expression levels of B16 *Csf1r*^+^ *Mafb*^+^ cluster-specific gene signatures (Fig. 1E) displayed on *t*-SNE plots of *Csf1r*^+^ *Mafb*^+^ clusters from PyMT tumors (Fig. 3C). Cells with top 70 percentile of signature expression level labeled in red. **(D)** Heatmap displaying expression levels of B16 *Csf1r*^+^ *Mafb*^+^ cluster-specific genes (Fig. 1E) by PyMT-infiltrating *Csf1r*^+^ *Mafb*^+^ cells (Fig. 3C). **(E)** Expression of *Il7r* (**left**) and *Vcam1* (**right**) by cells in PyMT *Csf1r*^+^ *Mafb*^+^ clusters (Fig. 3C). **(F)** Heatmap displaying expression levels of glycolysis- and oxidative phosphorylation (“OxPhos”)-associated genes in PyMT *Csf1r*^+^ *Mafb*^+^ cluster cells (Fig. 3C).

**Supplementary Figure 4.**
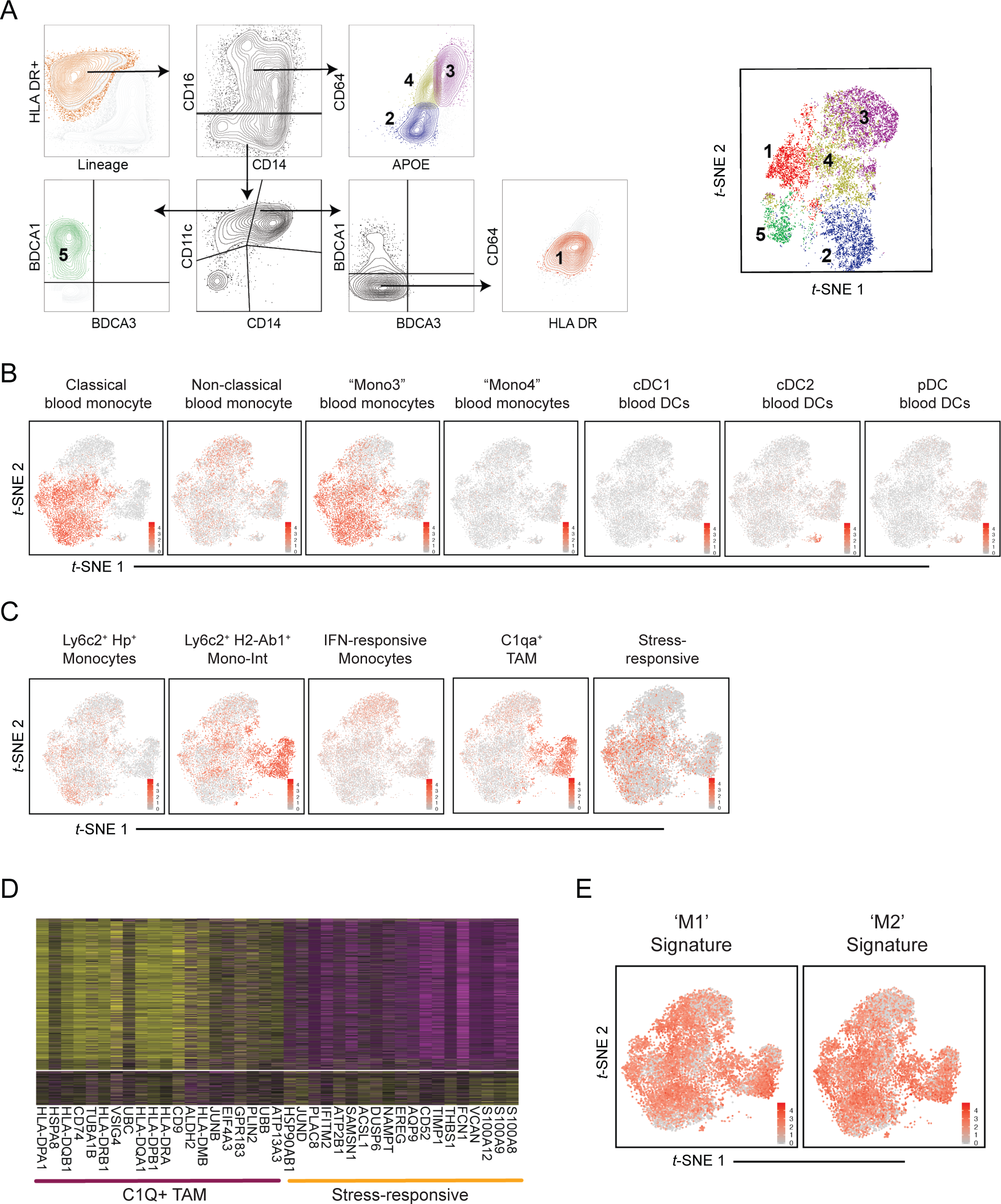
(A) Gating approach (**left**) and *t*-SNE plot (**right**) generated from flow cytometry analysis of the human RCC myeloid compartment. Example representative of more than 5 experiments. **(B)** Expression levels of previously established blood myeloid cell signatures^59^ displayed on *t*-SNE plot of myeloid cells from human RCC biopsy (Fig. 4B). **(C)** Expression levels of mouse B16 tumor myeloid cell signatures (Fig. 1E) displayed on *t*- SNE plot of *CSF1R*^+^ *MAFB*^+^ clusters from human RCC biopsy (Fig. 4C). **(D)** Heatmap of top 20 DE genes between human RCC stress-responsive (Cluster 4) and *C1Q*^+^ TAM (Cluster 3) clusters (Fig. 4B). Genes ranked by fold change. **(E)** Expression levels of “M1” (**left**) and “M2” (**right**) gene signatures (Fig. 3) displayed on *t*- SNE plot of *CSF1R*^+^ *MAFB*^+^ clusters from human RCC biopsy (Fig. 4C).

**Supplementary Figure 5.**
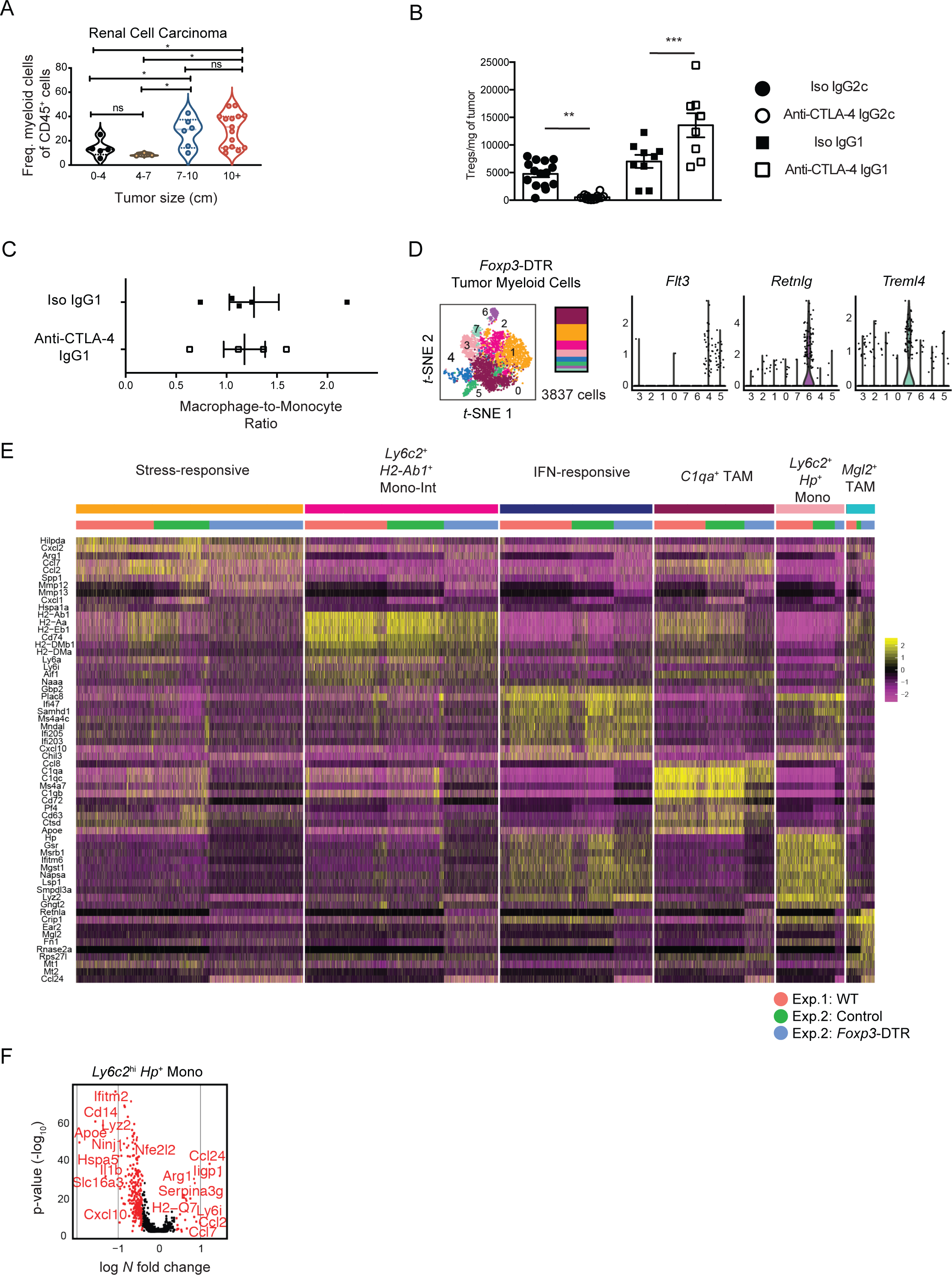
(A) Quantification by flow cytometry of myeloid cell frequency within CD45^+^ cells from human RCC biopsies. **(B)** Quantification of Treg cell number per milligram of B16 tumor from wildtype mice that were treated with anti-CTLA-4 antibodies (IgG2c or IgG1 clone) or corresponding isotype antibody controls (mean ± SEM). Data was pooled from two independent experiments. **(C)** Quantification of the ratio between macrophages (Ly6C^-^ F4/80^+^ CD64^+^) and monocytes (Ly6C^+^ CD11b^+^) in B16 tumors from wild-type mice treated with non-depleting anti-CTLA-4 (IgG1 clone) or isotype antibody control. Data is representative of two independent experiments. **p <0.01, ***p<0.001. **(D)** *t*-SNE plot of graph-based clustering of (**left**) and expression levels of select genes (**right**) of B16 tumor myeloid cells sorted from DT-treated *FoxP3*-DTR mice. **(E)** Heatmap displaying expression of top 10 DE genes between B16-infiltrating *Csf1r*^+^ *Mafb*^+^ clusters aggregated from wildtype (Fig. 1C), DT-treated control and DT-treated *FoxP3*-DTR mice. Genes are ranked by fold change. **(F)** Volcano plot displaying DE genes between B16 tumor *Ly6c2*^+^ *Hp*^+^ monocytes from DT-treated control and *FoxP3*-DTR mice. Genes with > 0.4 log-fold changes and an adjusted p value of 0.05 (based on Bonferroni correction), are highlighted in red. Genes of interest labeled.

**Supplementary Figure 6.**
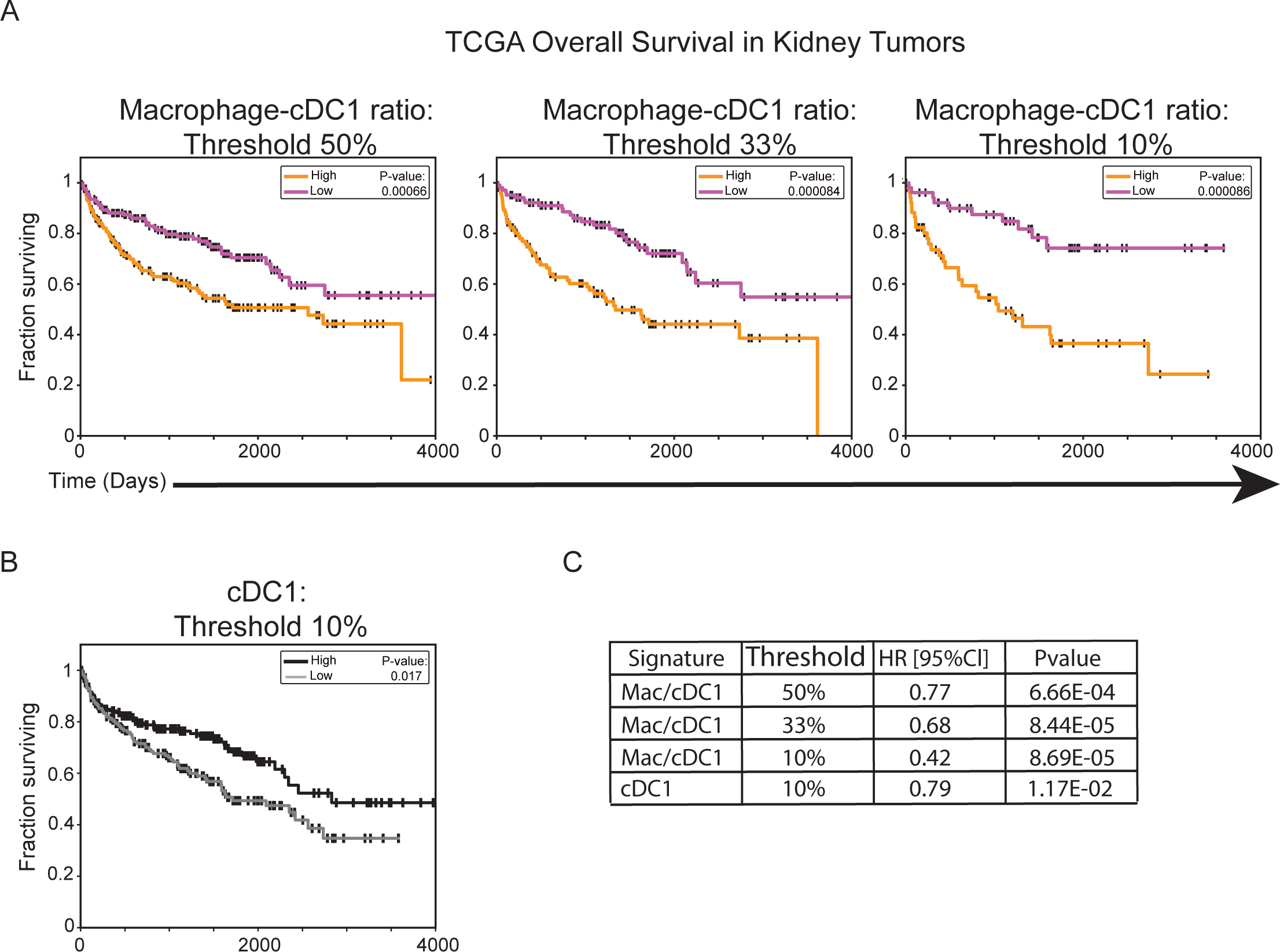
(A) Survival curves of kidney tumor patients whose TCGA tumor samples exhibited differential levels of the ratio of macrophage (Fig. 4E) to cDC1^53^ gene signature scores. A score threshold of 50% (l**eft**), 33% (**middle**), and 10% (**right**) was used to determine high and low comparison groups. **(B)** Survival curves of kidney tumor patients whose TCGA tumor samples exhibited differential levels of the cDC1 gene signature expression score^53^. A score threshold of 10% was used for patient group selection. **(C)** Summary of hazard ratios and p-values from analyses of kidney tumor samples from TCGA (**A,B**).

**Table S1. DE genes between B16 tumor myeloid populations.** DE analysis was performed between B16 tumor clusters (**Fig. 1B**). DE genes with log N fold change >0.5 and min.pct of 0.25 are listed.

**Table S2. DE genes between human RCC myeloid populations.** DE analysis was performed on RCC myeloid clusters (**Fig. 4B**), and DE genes with log N fold change >0.4 and min.pct of 0.25 are listed.

**Table S3. DE genes between monocyte/macrophage populations in control and Foxp3- DTR tumors.** DE analysis was performed on aggregated myeloid cell clusters from control and *Foxp3*-DTR (**Fig. 5D**). DE genes with log *N* fold change > 0.001 and min.pct of 0.1 listed. DE analysis for each cell cluster is listed on the specified tab.

